# Loss of age-associated increase in m^6^A-modified RNA contributes to GABAergic dysregulation in Alzheimer’s disease

**DOI:** 10.1101/2025.05.02.651974

**Authors:** Jenna L. Libera, Jacqueline T. M. Aw, Junming Hu, Tuyet-Anh Nguyen, Zihan Wang, Sophie J. F. van der Spek, Kamryn Schult, Luke Dorrian, Jordan Majka, Katarnut Tobunluepop, Sambhavi Puri, Alexander Knyshov, Juao-Guilherme Rosa, Yuran Ma, Nicholas M. Kanaan, Peter T. Nelson, Kate Meyer, Lei Hou, Julia TCW, Xiaoling Zhang, Benjamin Wolozin

## Abstract

**INTRODUCTION:** Dysregulated RNA metabolism is a significant feature of Alzheimer’s disease (AD), yet how post-transcriptional RNA modifications like *N*^6^-methyladenosine (m^6^A) are altered in AD is unknown.

**METHODS:** Here, we performed <u>d</u>eamination <u>a</u>djacent to <u>R</u>NA modification targets (DART-seq) on human dorsolateral prefrontal cortices to assess changes in m^6^A with nucleotide resolution.

**RESULTS:** In non-AD brains, m^6^A sites increased with age, predominantly within the 3′UTR of transcripts encoding tripartite synapse proteins. In contrast, AD brains lost the age-associated m^6^A site increase and exhibited hypomethylation of transcripts overall, including *MAPT*. Hypomethylated genes were involved with GABAergic signaling, glutamate transport, and ubiquitin-mediated proteolysis exhibited reduced expression, connecting m^6^A to synaptic excitotoxicity and disrupted proteostasis in AD. Site-specific m^6^A levels were linked with *GABRA1* expression and protein levels, but this relationship was abolished in AD.

**DISCUSSION:** Our findings provide insight into post-transcriptional features of dysregulated RNA metabolism in AD that are related to aging and GABAergic regulation.

**Graphical Abstract:** 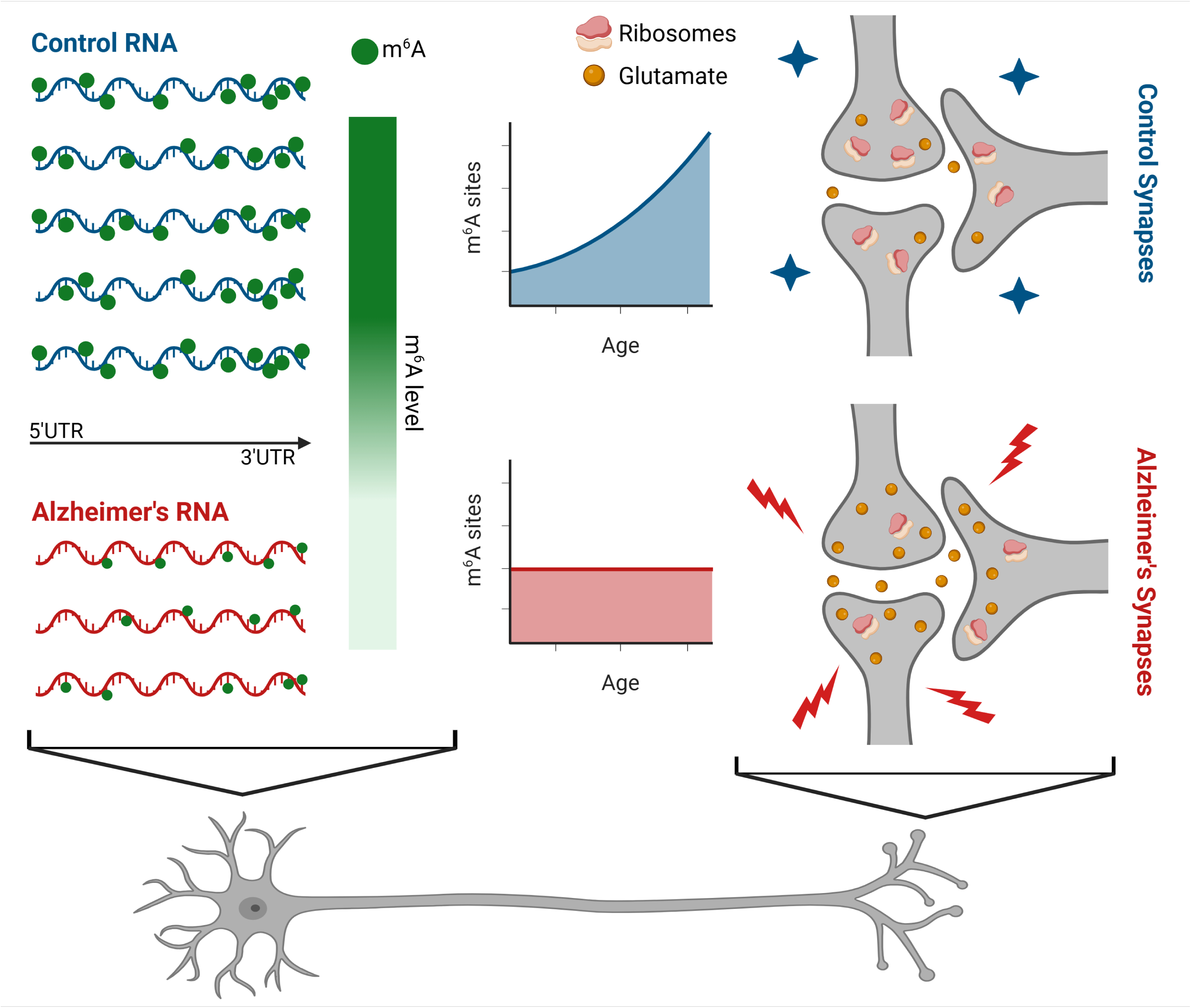

## 1 BACKGROUND

Alzheimer’s disease (AD) and other neurodegenerative disorders are increasingly linked to dysfunction in RNA metabolism. This dysfunction arises in part from the biochemistry of RNA binding proteins (RBPs) which form biological condensates that, if persistent, generate pathological aggregates. For example, in amyotrophic lateral sclerosis (ALS), TDP-43 mislocalization and aggregation drive cryptic splicing, resulting in production of abnormally spliced RNA and loss of the cognate mRNA.^1,2^ RNA-processing defects are also seen with other disease-linked RBPs, including HNRNPA2B1, FUS, Ataxin-2, TIA-1, and Musashi.^3^ In AD, stress-mediated phosphorylation stimulates oligomerization of microtubule-associated protein tau (MAPT) that in turn promotes its association with RBPs in biological condensates, such as stress granules.^4–7^ We reported that one of the strongest RBPs interacting with oligomeric tau is HNRNPA2B1, which regulates RNA metabolism via its binding to *N*^6^-methyladenosine (m^6^A) modified mRNA, one of the most abundant mRNA modifications.^8,9^ This finding raises the possibility that the m^6^A-transcriptome is altered in AD.

m^6^A is a dynamic RNA modification installed by methyltransferases (i.e., writers) and removed by demethylases (i.e., erasers).^10^ It is enriched in DRACH motifs (D = A/G/U, R = A/G, A = A or m^6^A if methylated, C = C, and H = A/C/U), and regulates diverse aspects of RNA metabolism, including transcription, splicing, nucleocytoplasmic export, localization, translation, stability, and stress granule formation.^11–13^ These functions are mediated by “readers”, or RBPs, that bind m^6^A-modified RNA.^14^ m^6^A readers include TDP-43, HNRNPA2B1 and FMRP, all relevant to neurodegeneration.^9,15,16^

Neurons uniquely leverage m^6^A for spatially localized translation in the axon, dendrites, and synapses particularly through 3’UTR-specific m^6^A labeling, and the synaptic m^6^A epitranscriptome has been implicated in neurodevelopmental and neuropsychiatric diseases.^17–19^ However, most existing studies rely on m^6^A-antibody-based approaches by methylated RNA immunoprecipitation and sequencing (MeRIP-seq), which lacks nucleotide-specific resolution.^18,20^ m^6^A antibody-based methods also show cross-reactivity by binding the m^6^Am RNA modification.^11^ To overcome these limitations, m^6^A sequencing method, termed “deamination adjacent to RNA modification targets” (DART-seq) was developed to enable single-nucleotide m^6^A mapping.^21^ This method uses a fusion protein of the m^6^A-binding YTH domain and the C-to-U deaminase APOBEC1 to achieve targeted editing of cytidines that follow methylated DRACH sites.^11^

We performed DART-seq on mRNA from the dorsolateral prefrontal cortex and report the first single-nucleotide characterization of m^6^A in the post-mortem human brain. and compared. When comparing AD and non-AD Control cases (n=19/cohort), we observe a striking age-dependent increase in m^6^A-modified sites in Controls, both globally and in genes associated with the tripartite synapse. Genes related to GABAergic signaling and glutamate transport show coordinated changes in methylation and transcript levels. In contrast, AD cases lack this age-dependent increase of m^6^A-modified sites and show hypomethylation of genes, including disease-linked genes like *MAPT*. Tripartite synapse genes *GABRA1, GABRA2, GABRA4,* and *SLC1A2* have hypomethylated 3’UTRs and reduced expression in AD, implicating excitotoxicity. Methylation level of specific 3’UTR sites in *GABRA1* is linked with its transcript and protein level in Control cases, but not in AD. These results identify m^6^A as a regulator of brain aging and that its dysregulation contributes to synaptic dysfunction in AD.

## 2 METHODS

### 2.1 Tissue samples

Brain tissues were obtained from the University of Kentucky Alzheimer’s Disease Research Center (UK-ADRC) at the Sanders Brown Center on Aging. Methods for recruitment and pathological workup of these cases have been described previously.^22,23^ To capture advanced progression of AD and provide clearer distinction to non-AD cases, dorsolateral prefrontal cortex (Broadman area 9) samples from Control and AD cases using pathological criteria for group designation (n=19/group) were used in this study.^24^ The demographics, clinical, neuropathological characteristics, co-pathology observations, and post-mortem intervals (PMI) of the cases are presented in Table S1. The Control and AD cohorts were matched for age, sex, PMI, and RNA quality.

### 2.2 Tissue homogenization and RNA extraction

Brain tissue was cut into ∼50mg pieces on dry ice. The tissue was then wrapped in tin foil, placed over a metal block half submerged in liquid nitrogen, pulverized by a hammer, and placed into a lysing tube (Fisher Scientific, #MP116913100). 1 mL of Qiazol (Qiagen, #79306) was added to each tube. Tissue was homogenized by a bead mill (Fisherbrand™, #15-340-163) for 40 s at 4 °C and then sat at room temperature for 5 minutes. 200 μL of chloroform was then added, tubes were shaken vigorously for 60 s, and then rested again at room temperature for 3 minutes. Samples were centrifuged for 15 minutes at 4 °C for 12,000 x g to separate the organic, middle, and aqueous phase. The aqueous phase was carefully transferred to a gDNA eliminator column (Qiagen, part of RNeasy Plus kit) and spun at 8000 x g for 15 s. The spun phase was then mixed with 70% EtOH and transferred to a spin column. The RNA purification steps were subsequently carried out as described using the RNeasy Plus kit with DNase I treatment (Qiagen, #74136). After elution with 40 µL RNase-free water, RNA was quantified and assessed for quality on the Agilent TapeStation 4200 using the RNA ScreenTape assay and for purity on the Nanodrop.

### 2.3 DART enzyme preparation and quantification

DART enzyme sequences (pCMV-APOBEC1-YTH [Addgene, #131636] and pCMV-APOBEC1-YTH^mut^ [Addgene, #131637]) were cloned into the pET-His6-MBP vector (Addgene, #29656, a gift from Kate Meyer). A glycerol stab of Rosetta^TM^ 2 (DE3) pLysS Singles^TM^ Competent Cells (Sigma Aldrich, #71402) was transformed with DART-enzyme expression vectors and grown in Terrific Broth (Thermo Fisher, #22711022, supplemented with 50 µg/mL Kanamycin) overnight at 37 °C with shaking at 200 rpm. The following day, 1 mL of starter culture was added to 90 mL of Boca Scientific Inc. Terrific Broth Auto-Induction Medium (Fisher Scientific, #NC1894479, supplemented with 50 µg/mL Kanamycin) and incubated for 7 hours at 37 °C with shaking at 200 rpm. Then, the cells were placed at 18 °C overnight with shaking at 200 rpm. Finally, the bacteria cultures were spun down at 6000 x g for 10 minutes at 4 °C, and the cell pellet was stored at -80°C until further use. The cell pellet was lysed using the EasyPrep Bacterial Protein Extraction Kit (Cepham, #10450). After thawing on ice for 30 minutes, 10 mL of lysis buffer (0.2% lysozyme, 2% nuclease) was added, and cells were homogenized and incubated for 30 minutes with mixing every 5-10 minutes. Cell debris was spun down at 12,500 x g for 1 hour at 4 °C to obtain a cleared protein extract. The protein extract was purified using the HiTrap Chelating columns (Cytiva Life Sciences, #17040901) on the Äkta pure^TM^ chromatography system (Cytiva Life Sciences). All steps were carried out using DEPC-treated water. HiTrap Chelating columns were first flushed with water, charged with 0.1 M cobalt(II) chloride, equilibrated with an equilibrium buffer (50 mM Na2HPO4, 100 mM NaCl, 10 mM imidazole, pH 7.4), and loaded with protein extract by a circulating pump following the manufacturer’s instructions. Next, the columns were washed with wash buffer (10 mM Hepes, 300 mM NaCl, 10 mM imidazole, pH 7.4), and the enzymes were eluted with a linear gradient from 15% to 55% of elution buffer (10 mM Hepes, 1 M NaCl, 300 mM imidazole, pH 7.4) using the Äkta pureTM system. Finally, purified enzyme was dialyzed using Slide-A-Lyzer Dialysis Cassettes (20,000 MWCO; ThermoScientific, #66012) in dialysis buffer (10 mM Tris-HCl, 100 mM NaCl, DTT, pH 7.4), mixed with glycerol to obtain a 20% glycerol solution, snapped frozen, and stored at -80 °C until further use. Protein purity and concentration were assessed against a BSA standard curve using SDS-PAGE followed by Coomassie stain.

### 2.4 DART reaction and RNA repurification

DART reactions were performed as previously described.^25^ A 50 µL reaction was set up in a 200 µL PCR tube (5 μL 10x DART buffer [100 mM Tris-HCL (pH 7.4), 500 mM KCl, 1 μM ZnCl_2_], 1 μL RNasin® Ribonuclease Inhibitor [40 U/µL, Promega, #N2615], 50 ng of total purified RNA, 250 ng of purified DART protein [either APOBEC1-YTH or APOBEC1-YTH^mut^], molecular grade RNase-free water to 50 µL). Tubes were gently mixed by tapping, briefly spun, and added to a pre-heated thermal cycler at 37 °C for 4 hours. After incubation, RNA was purified using the RNeasy Micro Kit (Qiagen, #74004) according to the manufacturer’s instructions and eluted with 14 µL of molecular grade RNase-free water. RNA was quantified and assessed for quality on the Agilent TapeStation 4200 using the High Sensitivity RNA ScreenTape assay.

### 2.5 cDNA synthesis, DART PCR, and editing quantification from Sanger sequencing

cDNA was generated with the SuperScript III Reverse Transcriptase kit (Invitrogen, #18080044) using 3 ng of repurified DART-enzyme-treated total RNA as input and 50 ng of random hexamer primers for amplification (Thermo Scientific, #N8080127). PCR was performed with AmpliTaq Gold® 360 Master Mix (Applied Biosystems, #4398881) using 1 µL of cDNA and 1.0 µM of forward and reverse gene-specific primers (list below). PCR products were sent to Quintara Biosciences for purification and Sanger sequencing. Percent C>U conversion was quantified using EditR.^26^

### 2.6 DART PCR primers

The *ACTB* primers were from a previous study.^27^ The *GABRA1* 3’UTR primers were designed in 4 sets to capture the 7 detected m^6^A sites by DART-seq (**see Table S2**).

### 2.7 cDNA library synthesis and sequencing processing

3 ng of total RNA was amplified to create cDNA libraries using the SMARTer® Stranded Total RNA-Seq Kit v2 (Takara Bio, #634411) following the manufacturer’s protocol. Libraries were generated for all n=38 APOBEC1-YTH-treated samples, while only a half of the libraries were generated for APOBEC1-YTH^mut^-treated samples: C5, C7, C8, C10, C11, C12, C15-C19, A4-6, A8, A10-13, A15, A16, and A18 (***see Table S1***). cDNA library quality, appropriate fragment size (300-400 bp), and concentration were determined using the Agilent TapeStation 4200 and D1000 ScreenTape Assay. The cDNA libraries were normalized and pooled at a concentration that achieved >20,000,000 unique reads per sample and run on the Illumina Next-seq 2000 (100 cycles with 50 x 50 paired ends) by the Microarray and Sequencing Resources Facility at Boston University Medical Center to generate 1.2 billion total reads.

Fastq files from the DART reactions were obtained from BaseSpace Illumina. Quality control (QC) of the sequence data was performed using FastQC (v0.12.1) which assessed read quality, GC content, duplication levels, and adaptor contamination.^28^ Reads passing initial QC were aligned to the human reference genome (GRCh38.111) using STAR (version 2.7.9a). This mapped sequencing reads to the reference genome using single-pass alignment, generating sorted BAM files for downstream analysis.^29^ The resulting BAM files for each sample contained mapped paired-end reads and a corresponding alignment report file. Gene and isoform levels were quantified using RSEM (version 1.3.1) and Bowtie2 (version 2.3.4.1) and then annotated using Homo sapiens GRCh38.111.gtf annotation files.^30,31^ For each sample, this process generated gene-level files containing gene ID, length, effective length, expected count, CPM, and FPKM, which were then combined across samples based on gene IDs for downstream analysis. ‘Batch effects in the quantified gene expression data were assessed using principal component analysis (PCA).

### 2.8 m^6^A site identification by C>U editing and visualization

The Bullseye pipeline was used to identify m^6^A-modified sites (i.e., C>U editing).^17,25,27,32^ First, the parseBAM.pl script was used to parse the sequences from the coordinate-sorted BAM files. Then, a matrix containing the number of A, T, G, C, and N at all positions with a minimum coverage of 10 reads was created for each APOBEC1-YTH-treated sample. For APOBEC1-YTH^mut^-treated samples, all BAM files were merged using Samtools and parsed as a reference matrix. The Find_edit_site.pl script was then used to find C>U mutations by comparing each APOBEC1-YTH sample to the APOBEC1-YTH^mut^ reference matrix. We defined true, high-confident m^6^A sites for each sample if a minimum number of C>U mutation events at a particular site was 2, the site had an editing ratio (i.e., C>U site counts/total site counts) between 0.10-0.95, and the APOBEC1-YTH editing ratio enrichment over APOBEC1-YTH^mut^ editing ratio was at least 1.5 times higher. Finally, sense C>U (and antisense G>A) transitions were filtered for presence of the surrounding DRACH motif using custom scripts and the GenomicRanges (10.18129/B9.bioc.GenomicRanges), BSgenome (10.18129/B9.bioc.BSgenome), Biostrings (10.18129/B9.bioc.Biostrings), and gUtils R packages.^33,34^

Genomic features corresponding to m^6^A modifications were first imported from a BED file generated using Bullseye results, employing the Guitar package.^35^ A comprehensive transcriptome annotation for Homo sapiens (GRCh38.111) was generated from a GTF file via the makeTxDbFromGFF function. The spatial distribution of m^6^A sites along mRNA transcripts was subsequently visualized using the GuitarPlot function. This approach mapped the input BED file to transcriptomic landmarks with a focus on both 5′ and 3′ regions, as indicated by the headOrtail parameter, and normalized the data by scaling each m^6^A site to a relative position (0– 1 range) within transcript regions to account for region length, and by applying site-specific weights based on transcript-level overlap and ambiguity to ensure accurate regional enrichment. Custom filtering (mapFilterTranscript = TRUE) was applied to ensure precise localization of features, and the plot aesthetics were refined to enhance the interpretability of the resulting visualization.

For enrichment analysis of DRACH motif, background sequences were generated by shuffling the input BED coordinates and extracting both target and background sequences from the human genome (GRCh38.111) using bedtools. HOMER tools were then applied—using findMotifs.pl for motif detection and annotatePeaks.pl with findMotifsGenome.pl for peak annotation—to identify enriched customized motifs in the target regions relative to the background.^36^

### 2.9 Gene expression analysis and visualization

Differential expression analysis was conducted using the DESeq2 package in R.^37^ Initially, raw count data were imported and subjected to a filtering step wherein only those genes exhibiting counts ≥10 in at least 19 samples were retained. Count normalization was subsequently performed using counts per million (CPM) to adjust for library size differences. To mitigate the influence of confounding variables, covariates including pre-extraction RNA integrity number (pre_dart_RIN), age, sex, and post-mortem interval (PMI) were incorporated into the model. The final design formula was specified as ∼ pre_dart_RIN + Age + sex_num + PMI + AD_Ctr, with AD_Ctr representing the binary classification of AD versus Control. Differentially expressed genes (DEGs) between the AD and Control groups were determined using the Wald test; DEGs with an adjusted p-value (padj) ≤ 0.05 were deemed statistically significant.

To evaluate the similarity between our data and publicly available MSBB datasets, we computed Spearman correlations between the mean gene expression levels in our dataset and those across the BM 10 MSBB dataset (https://adknowledgeportal.org/). The MSSB data were generated from post-mortem brain tissue collected through the Mount Sinai VA Medical Center Brain Bank and were provided by Dr. Eric Schadt from Mount Sinai School of Medicine. We first calculated log2 counts per million (log2 CPM) using EdgeR, identified shared genes, extracted their log-transformed expression values, and performed correlation analysis.^38^ The results, including correlation coefficients and p-values, were visualized using scatter plots with fitted regression lines.

PCA and volcano plots for DMGs and DEGs were generated in R using the ggplot2 package.^39^ KEGG pathway analysis was performed using ShinyGo.^40–42^ GSEA was performed and visualized with dot plots and a network plot in R using the clusterProfiler package.^43,44^ Heatmaps for DAS and genes with overlapping DART-seq and RNA-seq GSEA terms were generated in R using the ComplexHeatmap package.^45^

### 2.10 Synaptosome isolation, BCA assay, and protein normalization

Synaptosome isolation was performed using Syn-PER™ Synaptic Protein Extraction Reagent (Thermo Scientific, #87793) following the manufacturer’s protocol. Briefly, ∼50 mg of brain tissue was dounced in Syn-PER Reagent supplemented with 1x of Halt™ Protease and Phosphatase Inhibitor Cocktails (100x, Thermo Scientific, #78440) on ice with ∼10 slow strokes. The homogenate was transferred to a 1.7 mL microcentrifuge tube and spun at 1200 x g for 10 minutes at 4 °C. Supernatant was collected and the spin was repeated. Then, 20% of the supernatant was saved as the homogenate fraction and the other 80% was placed in a new 1.7mL tube and centrifuged at 15,000 x g for 20 minutes at 4 °C. The supernatant (i.e., cytosolic fraction) was saved and the pellet containing the synaptosomal fraction was resuspended in Syn-PER at 20% of the original volume used. Then, the fractions were quantified using the Pierce™ BCA Protein Assay Kit (Thermo Scientific, #23225 and #23227) according to the manufacturer’s protocol. Absorbance was measured using the SpectraMax i3x Platform. Protein stocks were normalized across all samples to achieve the same concentration and volume after adding 1x Bolt™ Sample Reducing Agent (10x, Invitrogen, #B0009) and 1x of Bolt™ LDS Sample Buffer (4x, Invitrogen, #B0007). Finally, samples were boiled for 10 minutes at 95 °C and stored at -20 °C until use.

### 2.11 Immunoblot

Homogenate and synaptosomal fractions (3-10 µg, see Table S3) were loaded into a 4-15% Criterion^TM^ TGX Stain-Free^TM^ Protein Gel (Bio-Rad, #5678085) and were run using 1x Bolt MES SDS Running Buffer (20x, Life Technologies, #B0002). Afterwards, the gel was scanned using the Bio-Rad ChemiDoc^TM^ XRS+ imager to obtain total protein bands for densitometry normalization. The gel was transferred to a PDVF membrane using the iBlot 2 System (Thermo Fisher Scientific) template #1. Once transferred, the membrane was blocked in Tris-buffered saline + 0.05% Tween (TBST) with 5% non-fat dry milk for 60 minutes shaking gently at room temperature. The membrane was then incubated in primary antibody overnight (list below) shaking gently at 4 °C in a solution of either TBST +1% BSA or phosphate buffered saline + 0.05% Tween (PBST) + 1% BSA (see Table S3). After 24 hours, the membranes were washed 3 times in TBST or PBST for 5 minutes each, shaking gently at room temperature. Then, the membranes were incubated in secondary antibody in TBST or PBST for 60 minutes shaking gently at room temperature. Secondary antibody solution was removed and membranes were washed 3 more times in TBST or PBST for 5 minutes each with gentle shaking at room temperature. Afterwards, membranes were incubated for ∼15-30 seconds in Super Signal West Pico PLUS stable peroxide and luminol/enhancer (Thermo Scientific, #34580) and then imaged using the Bio-Rad ChemiDoc^TM^ XRS+ system. Densitometry quantification of images was performed using ImageJ.

### 2.12 *GABRA1* qPCR

Following RNA extraction, 1 µg of total RNA was amplified into cDNA using the High-Capacity cDNA Reverse Transcription Kit (Thermo Fisher Scientific, #4368814). cDNA was prepared for qPCR using Ssoadvanced™ Universal SYBR Green (Bio-Rad #1725270) and gene-specific primers (list below). Ct levels of *GABRA1* were normalized to Ct levels of *GAPDH* and fold change was determined using the 2^-ΔΔ*CT*^ method.^46^

### 2.13 qPCR primers

See Table S4.

### 2.14 Statistical analysis and figure generation

Statistical analyses were performed in GraphPad/Prism 10.1.2. Anderson-Darling, D’Agostino, Shapiro-Wilk, and Kolmogorov-Smirnov tests for normality were performed on all comparisons. If at least half of the tests were significant, the data were considered normally distributed. Unpaired t-tests were used for comparing normally distributed means of 2 groups. Otherwise, Mann-Whitney tests were performed. One-way or two-way ANOVAs, followed by Tukey’s HSD post hoc analyses, were used for comparing normally distributed means of 3 or 3+ groups. Otherwise, Kruskal-Wallis tests followed by Tukey’s HSD post hoc analyses were performed. For all tests, p-value was set to 0.05 and mean ± standard deviation was reported. Results were visualized in GraphPad/Prism 10.1.2 or in R. Workflow illustrations were made with Biorender.

## 3 RESULTS

### 3.1 Validation of DART-seq confirms detection of m^6^A sites in human post-mortem brains

Detection of m^6^A sites at the single-nucleotide level was accomplished by performing DART-seq on RNA from human post-mortem brains (**Table S1**). We analyzed age-matched (**Fig. S1A**) and sex-matched (**Fig. S1B**) non-AD Control and AD brains (n=19 each). As per the diagnostic criteria, AD cases had higher tau burden (**Fig. S1C**) and amyloid burden (**Fig. S1D**) than Control cases. The majority of AD cases were *APOE* ε*3/*ε*4* carriers while most Control cases were *APOE* ε*3/*ε*3* carriers (**Fig. S1E**). Notably, in both cohorts, brain samples were collected with a mean post-mortem interval of ∼2.5 hours (**Fig. S1F**); the RNA quality for each sample was >7 RIN^e^ pre-DART reaction and >6 RIN^e^ post-DART reaction (**Fig. S1G**), highlighting robust sample quality.

To ensure feasibility of the DART-seq method on our human post-mortem brains, we standardized a workflow involving automated sample homogenization, multiple rounds of RNA purification, and quality control checks of RNA and cDNA libraries using a Tapestation 4200 (Agilent, **Fig. S2A**). Two m^6^A sites along the *ACTB* transcript, A1222 and A1248, were previously reported to have high levels of m^6^A.^47^ We validated m^6^A level of these sites in a subset of the samples (n=3/cohort) by incubating RNA in parallel conditions with APOBEC1-YTH to induce C>U editing and with APOBEC1-YTH^mut^ as a negative control, performing PCR to amplify the sites, and quantifying the proportion of C>U editing as given by Sanger sequencing (**Fig. S2B**). DART-seq on our full sample set used next generation sequencing and the Bullseye pipeline to uncover novel m^6^A sites (**Fig. 1A**).^25,27,32,48^ Indeed, our C>U editing was ∼16% at A1222 and ∼32% at A1248 of RNA incubated with APOBEC1-YTH; RNA that was incubated with APOBEC1-YTH^mut^ was edited ∼50% less (**Fig. S2C**). These levels match prior reports.^47^ We also profiled C>U conversion of A1035, a non-methylated DRACH motif, and observed almost no editing (∼1-5%, **Fig. S2B**). This confirms that DART is specific to m^6^A-occupied DRACH motifs as previously reported.^21,27^ Importantly, DART-seq identified sites A1222 and A1248 across all 38 samples, showing similar C>U conversion to our PCR validation (**Fig. S2D**). Together, these results demonstrate validity of DART-seq on human post-mortem brains.

**FIGURE 1:**
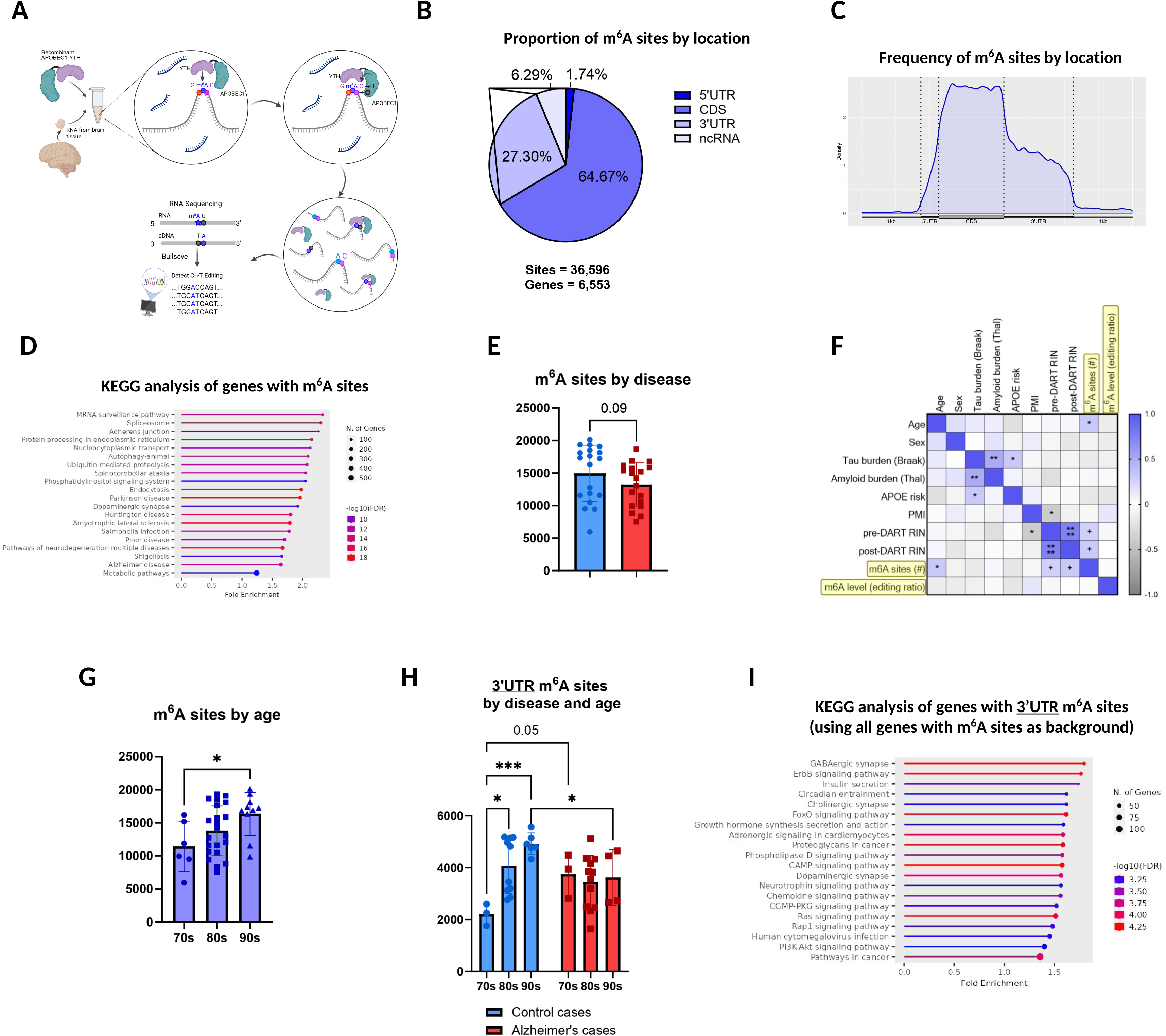
Alzheimer’s disease cases exhibit loss of age-dependent m^6^A increase in the 3’UTR. (A) Overview of DART reaction and Bullseye program. Purified RNA from brain tissue is incubated with recombinant APOBEC1-YTH (or APOBEC1-YTH^mut^ as a negative control). YTH binds to m^6^A modifications in a DRACH motif and APOBEC1 deaminates adjacent Cs into Us (YTH^mut^ exhibits reduced binding and therefore reduced deamination events by APOBEC1), which is read and quantified as C>T edits following reverse transcription, sequencing, and detection by Bullseye. (B) Pie chart showing percentage of m^6^A sites along different mRNA domains and on ncRNA. (C) Density plot showing m^6^A site distribution along mRNA position. (D) KEGG analysis of genes with m^6^A sites. (E) Comparison of m^6^A site quantification between Control (n=19) and AD (n=19) cases. *P* = 0.0850 by Mann-Whitney U test. Error bars represent mean ± standard deviation. (F) Correlation matrix of tissue sample and case demographic information with RNA quality and m^6^A site and level quantification of all 38 cases. Pearson correlation coefficients range from -1.0 to 1.0. ^+^ *P* < 0.10, * *P* < 0.05, ** *P* < 0.01 (G) Comparison of m^6^A site quantification of all cases grouped by age decades (70s, 80s, and 90s). * *P* < 0.05 by one-way ANOVA with Tukey’s HSD post-hoc test. Error bars represent mean ± standard deviation. (H) Comparison of 3’UTR-localized m^6^A site quantification between Control and AD cases grouped by age. (* < 0.05, *** < 0.001) *P* values were generated by two-way ANOVA with Tukey’s HSD post-hoc test. Error bars represent mean ± standard deviation. (I) KEGG analysis of genes with 3’UTR-localized m^6^A sites (n=2,968). All genes with m^6^A sites, regardless of site location, were used as a background reference (n=6,878).

### 3.2 Loss of age-dependent m^6^A sites on genes involved in synaptic function occurs in Alzheimer’s disease

DART-seq of all 38 brains revealed 36,596 m^6^A sites across 6,878 genes (**Fig. 1B, Table S5**). The distribution of m^6^A sites across the mRNA transcript showed that sites were most abundant in the CDS (**Fig. 1C**), matching distribution curves from prior DART-seq studies.^15,49^ These sites were significantly enriched for the DRACH motif (*P*=1.0e-331, **Fig. S3A**). A KEGG pathway analysis included RNA metabolism (e.g., “mRNA surveillance pathway”, “Spliceosome”, “Nucleocytoplasmic transport”) among the top 5 enriched groups of genes with m^6^A sites, reflecting standard m^6^A pathways, followed by many neurodegenerative terms including “Alzheimer disease” (**Fig. 1D**).

A trending decrease in the number of total m^6^A sites was observed in AD cases compared to Control cases (**Fig. 1E**). There were also trending decreases in the number of 5’UTR, CDS, 3’UTR, and ncRNA m^6^A sites for AD cases compared to Control cases (**Fig. S3B-E**). This prompted us to check whether differences in writer, reader, and eraser expression were present between Control and AD samples (**Fig. S3F-G**). Only *YTHDC2* and *FTO* were significantly downregulated, albeit at small fold changes. Since AD cases exhibit less m^6^A modifications overall, mild *FTO* downregulation could signal more complex m^6^A regulation involving other molecular mechanisms to result in the change. Given the heterogeneity of post-mortem human brain samples, we investigated whether patient characteristics and co-pathologies associated with changes in m^6^A. Here, we identified age as an endophenotype that correlated significantly with the total number of m^6^A sites (**Table S5; Fig. 1F**). Increasing age (beginning at age 70 years) associated with increased m^6^A sites of specific mRNA domains, including the 5’UTR, CDS, and 3’UTR, but not on ncRNA (**Fig. 1G, Fig. S4A-D**). Interestingly, the correlation of age with m^6^A sites was seen only in the Control cases; AD cases exhibited no correlation of age with the amount of m^6^A sites. The stable number of m^6^A sites in AD across age suggests that the disease process disrupts the typical increase in m^6^A site number with aging (**Fig. S4E**). The age-dependent increase in m^6^A sites was seen most strongly in the 3’UTR among Control cases but was also present in the 5’UTR, CDS, and ncRNAs (**Fig. 1H, Fig. S4F-H**).

To better understand the role of m^6^A in the 3’UTR, we performed a KEGG pathway analysis on genes with 3’UTR m^6^A sites across all cases, which revealed “GABAergic synapse” as the most highly enriched term (**Fig. 1I**). Gene ontology (GO) Cell Compartment (CC) analyses strengthened the link of m^6^A to synaptic biology, showing “Glutamatergic synapse” as the most highly enriched term among genes with 3’UTR-localized m^6^A sites (**Fig. S5A**). Synaptic terms were also among the most enriched in all m^6^A-containing genes and gene locations (except the CDS; **Fig. S5A-E**). These data suggest that increasing m^6^A-modified sites is a feature of neurologically normal aging, which fails to occur in AD. The most vulnerable sites and associated pathways affected by the disease occur at 3’UTR sites involved in GABAergic functioning and glutamatergic synapses.

### 3.3 Differentially abundant m^6^A sites converge on tau phosphorylation, RNA metabolism, and GABA-A receptors

Next, we looked at whether there were differential m^6^A sites between Control and AD cases that could be related to disease mechanisms and synaptic function. To achieve this, we analyzed the number and level of m^6^A methylation between Control and AD cases at each m^6^A site and determined whether that site was overrepresented among Control or AD cases. This was performed using three analyses, which identified differentially methylated sites (DMS, **Table S6**), differentially abundant sites (DAS, **Table S7**), and differentially methylated genes (DMG**, Table S8, Fig. S6**). Sites that were present in >50% of both Control and AD cases were subjected to differential methylation analysis. Sites were differentially abundant in Control cases if >50% of cases had a site and <10% of AD cases had a site. Conversely, DAS in AD cases had >50% representation and <10% representation in Control cases. Methylation level of all sites, regardless of case representation, were summed and averaged across an entire gene or domain of that gene (e.g., 5’UTR, CDS, 3’UTR, ncRNA) and subjected to differential analysis, a method previously described in Flamand & Meyer.^49^

There were 229 differentially hypomethylated sites in AD and 84 differentially hypermethylated sites in AD; both hypo- and hypermethylated DMS were found predominantly in the CDS region, followed by the 3’UTR, on ncRNA, and in the 5’UTR (**Fig. S7A-B**, **Table S6**). The 3’UTR contained the highest relative amount of hypermethylated to hypomethylated sites in AD, whereas the 5’UTR contained the lowest relative amount of hypermethylated to hypomethylated sites in AD (**Fig. S7C**). GO BP analyses revealed associations with neuronal and synaptic processes in both hypo- and hypermethylated DMS (**Fig. S7D-E)**, which we explored further in our DAS analysis.

Control cases displayed 328 DAS across 309 genes while AD cases had only 137 DAS across 131 genes (**Fig. S8A-B**). The proportion of DAS identified between Control and AD cases was similar among all gene domains. (**Fig. S8C**). Among the most relevant DAS were those found in genes involved with tau phosphorylation status (e.g., *FYN*, *PPP1CB*, *PPP2R5C*) tau misfolding (e.g., *HSP90* paralogs), stress surveillance (e.g., *EIF2AK2/PKR*, *HIF1A*) stress granules (e.g., *TAF15*, *HNRNPC*, *DDX5*, *CAPRIN1*, *G3BP1*), and glutamatergic/GABAergic receptors (e.g., *GRIN2A*, *GRIN2B*, *GABRA* paralogs) (**Fig. S8D**). In particular, the DAS on *FYN*, *HSP90AA1*, and *TAF15* were found uniquely in AD cases and were completely absent in Control cases (**Fig. S8E**), suggesting that these may be key m^6^A sites that mediate pathological processes among AD cases specifically. The link between DAS and disease was also evident with a GO CC analysis; phosphatase complexes were among the most enriched terms for DAS in AD cases (**Fig. S8F**). In contrast, DAS in Control cases showed the most enrichment among GABAergic processes, such as “Benzodiazepine receptor activity”, by GO Molecular Function (**Fig. S8FG**). Together, these data suggest that there are cohort-exclusive m^6^A sites that have unique, different functions in the non-AD brain (i.e., regulating GABAergic signaling) and the AD brain (i.e., regulating tau phosphatases).

### 3.4 m^6^A-modified transcripts in Alzheimer’s disease are mostly hypomethylated and associate with downregulated genes that encode synaptic proteins

Expanding our analysis from single sites to gene-wide methylation, we found that almost all DMGs were strikingly hypomethylated (n=340; n=26 hypermethylated) in AD (**Fig. 2A**, **Table S8**), while RNA-seq revealed a more equal number of upregulated (n=231) and downregulated (n=377) (adjusted p-value < 0.05) differentially expressed genes (DEGs, **Fig. 2B, Tables S9 & S10**). The RNA-seq is consistent with prior datasets, such as from the Mount Sinai Brain Bank (MSBB, https://adknowledgeportal.org/, **Fig. 2C**). The global reduction of m^6^A level among AD cases occurs independently of transcript level; little overlap was observed between DMGs and DEGs, suggesting that changes in m^6^A do not generally affect transcript abundance (**Fig. 2D**). Similarly, domain-specific methylation changes (i.e., mean levels of methylation that include all m^6^A sites within specific regions of the gene corresponding to the 5’UTR, CDS, 3’UTR, or ncRNAs) also show little overlap with DEGs (**Fig. S9A-H)**. Overall, these results reveal that transcripts are largely hypomethylated in AD and do not generally associate with site-specific m^6^A changes.

**FIGURE 2:**
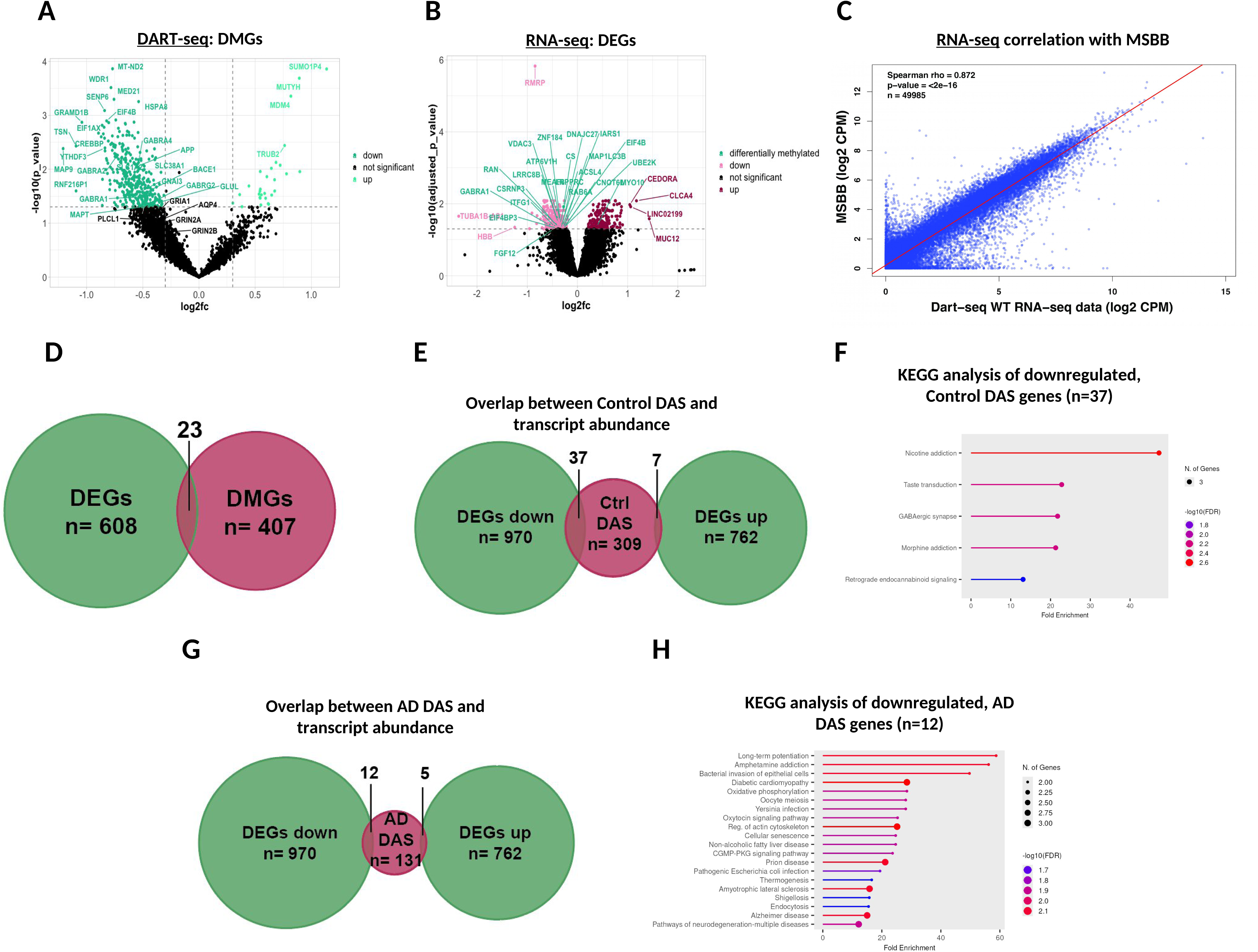
Globally hypomethylated transcripts in AD include *MAPT* and *APP* whose m^6^A level associates negatively with Braak stage. (A)Volcano plot of DART-seq revealing differentially methylated genes (DMGs). Y-axis represents –log10 p-value (dashed horizontal line is raw p = 0.05). X-axis represents log2 fold change with a cutoff of ± 0.3125 (dashed vertical lines). (B) Volcano plot of RNA-seq revealing differentially expressed genes (DEGs). Genes in green denote those that are also differentially methylated. Y-axis represents –log10 p-value (adjusted) with a significance cutoff (dashed horizontal line) of 1.3. X-axis represents log2 fold change. (C) Correlation of current RNA-seq dataset with Mount Sinai Brain Bank (MSBB) from brain region BM 10. Y-axis represents log2 counts per million (CPM) of the MSBB dataset. X-axis represents log2 CPM of the current RNA-seq dataset. (D) Euler plot showing the number of DEGs (n=609) and DMGs (n=407) and genes that are both differentially expressed and differentially methylated (n=23).

### 3.5 Characterization of tau spread via Braak staging reveals positive associations between amyloid burden and *APOE* genotype and a negative association with *MAPT* A5942 m^6^A level

A substantial hypomethylated DMG among AD cases is *MAPT*, the gene that encodes tau protein. *MAPT* is methylated exclusively in the 3’UTR and does not show differential mRNA expression (**Fig. S10A, see Table S10**). We proceeded to characterize MAPT given our prior studies identifying its relationship to m^6^A and the reader, HNRNPA2B1.^8^ To check for protein levels, we purified brain lysate from our Control and AD cases and quantified levels of MAPT by immunoblot. In AD cases, there was a 1.9x increase in MAPT compared to Control cases (**Fig. S10B-D**). Although an accumulation of aggregated MAPT in AD could account for this observed increase, these findings raise the possibility of a contribution of m^6^A-mediated regulation of protein synthesis, as this has been demonstrated before in cultured human and mouse neurons.^18,49,50^

We proceeded to explore the interaction between m^6^A and MAPT protein levels by examining AD endophenotypes. MAPT protein level positively associated with tau burden (i.e., Braak stage; **Fig. S10E**). DART-seq revealed two methylated DRACH sites at positions A4375 and A5942 of *MAPT* (**Fig. S10E, see Tables S5, S6**); the m^6^A level at site A5942 was negatively associated with Braak stage, suggesting that m^6^A level of this site may correspond to enhanced tau pathology. These findings reflect the possibility that the accumulation of tau signals back to RNA processing via m^6^A modification of *MAPT*. They also highlight differences of m^6^A regulation at the site-level and gene-level and reveal that methylation of *MAPT* m^6^A site A5942 is tightly linked to the clinical staging of tau spreading.

### 3.6 GABAergic synapses and proteolysis show strong gene set enrichment for m^6^A-linked expression

To determine concordant differences among groups of genes between Control and AD cases, we performed gene set enrichment analyses (GSEAs). The most enriched groups of the DART-seq dataset include many synaptic terms (e.g., “Synaptic vesicle cycle”, “GABAergic synapse”, and “Glutamatergic synapse”); other enriched groups include protein-related terms (e.g., “Ubiquitin mediated proteolysis”, “Protein processing in the endoplasmic reticulum”, and “Motor proteins”; **Fig. 3A, Fig. S11, Table S11**). Synaptic terms (e.g., “Synaptic vesicle cycle” and “GABAergic synapse”) are also enriched in the RNA-seq dataset with many more neurodegenerative terms included (e.g., “Parkinson disease”, “Prion disease”, and “Alzheimer disease”, **Fig. 3B, Table S12**). Other terms enriched in the RNA-seq dataset are related to proteostasis (e.g., “Ribosome”, “Proteasome”, and “Ubiquitin mediated proteolysis”). Importantly, four enriched gene sets overlapped between DART-seq and RNA-seq: “GABAergic synapse”, “Nicotine addiction”, “Synaptic vesicle cycle”, and “Ubiquitin mediated proteolysis” (**Fig. 3C-G, Fig. S11**). This suggests that these gene sets in particular have a strong link between differential m^6^A level and mRNA level in AD. The GABAergic synapse gene set includes multiple elements beyond the obvious signature of GABAergic receptors, including *SLC38A1* (i.e., *SNAT1*, a glutamine transporter) and *GLUL* (i.e., glutamine synthetase that removes glutamate by conversion to glutamine). The glutamate transporter *SLC1A2* (i.e., *EAAT2* or *GLT-1*) was also included in “Synaptic vesicle cycle”, linking potential impairment in glutamate uptake from synapses with excitotoxicity in AD. Other terms enriched in “Synaptic vesicle cycle” are *ATP6V* genes, *SNAP25*, and *CLTC* indicating the potential dysfunction of synaptic vesicle acidification, exocytosis, and endocytosis in AD (**Fig. 3F**). Neddylation and ubiquitination are also important elements of synaptic function (although not restricted to synaptic function) and correspondingly the term “Ubiquitin mediated proteolysis” showed enrichment for genes including *UBA*, *UBE*, and *CUL* paralogs (**Fig. 3G**). Together, these data suggest that in AD, hypomethylated genes contribute to aberrant functioning of GABAergic signaling, synaptic vesicle cycling, and ubiquitin-mediated proteolysis by associating with reduced mRNA expression.

**FIGURE 3:**
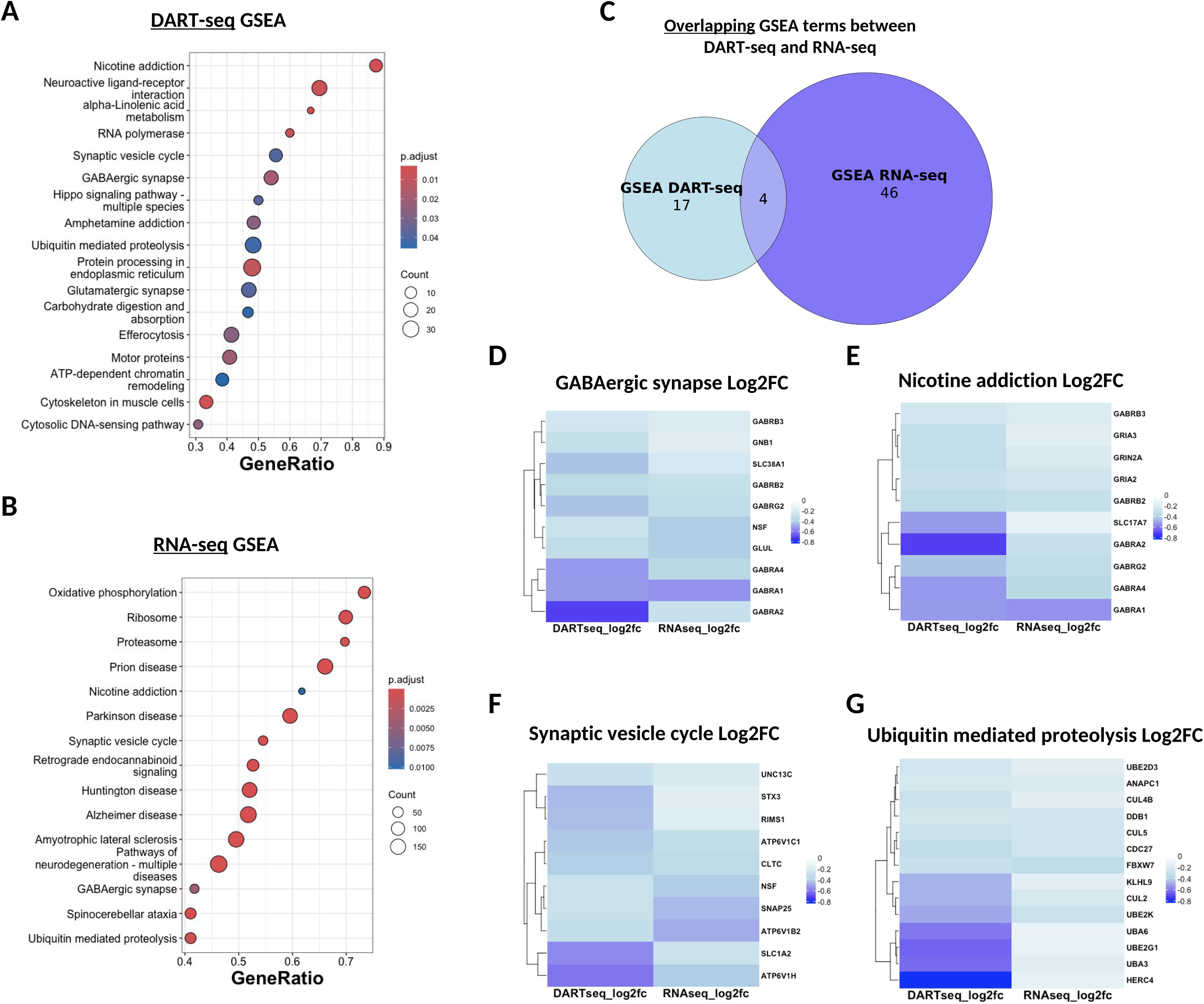
DART-seq and RNA-seq gene enrichment analyses reveal overlapping gene sets involved in GABAergic synapses, synaptic vesicle cycling, and proteolysis. (A) Dot plot of all GSEA terms from DART-seq depicting the significance (color), gene count (circle size), and gene ratio (x-axis) of each term. Complete GSEA analysis can be found in Table S10. (B) Dot plot of select GSEA terms from RNA-seq depicting the significance (color), gene count (circle size), and gene ratio (x-axis) of each term. Complete GSEA analysis can be found in Table S11. (C) Euler plot showing the number of GSEA terms from DART-seq (n=17) and the number of GSEA terms from RNA-seq (n=46) and terms that are shared between both GSEAs (n=4). (D) Heat map showing the log2 fold changes from DART-seq (left column) and RNA-seq (right column) of enriched genes (right labels) from the “GABAergic synapse” GSEA term. (E) Heat map showing the log2 fold changes from DART-seq (left column) and RNA-seq (right column) of enriched genes (right labels) from the “Nicotine addiction” GSEA term. (F) Heat map showing the log2 fold changes from DART-seq (left column) and RNA-seq (right column) of enriched genes (right labels) from the “Synaptic vesicle cycle” GSEA term. (G) Heat map showing the log2 fold changes from DART-seq (left column) and RNA-seq (right column) of enriched genes (right labels) from the “Ubiquitin mediated proteolysis” GSEA term.

### 3.7 Alzheimer’s disease brains lack an age-related m^6^A increase of tripartite synapse genes

We next determined which genes are shared among the enriched synaptic gene sets from DART- seq to identify synaptic targets linked to m^6^A-mediated synaptic impairment in AD. Our network plot revealed a striking coalescence of genes linked to the tripartite synapse, including GABAergic receptors (*GABRA1*, *GABRA2*, *GABRA4*), glutamatergic receptors (*GRIA1*, *GRIA2*, *GRIA3*, *GRIN2A*, *GRIN2B*) and astrocytic glutamate/glutamine interacting proteins (*SLC1A2*, *SLC1A3*, *SLC38A1*, *GLUL*), as well as signaling molecules regulating this axis (kinases, phosphatases, calcium binding proteins (**Fig. 4A, yellow circles**).

**FIGURE 4:**
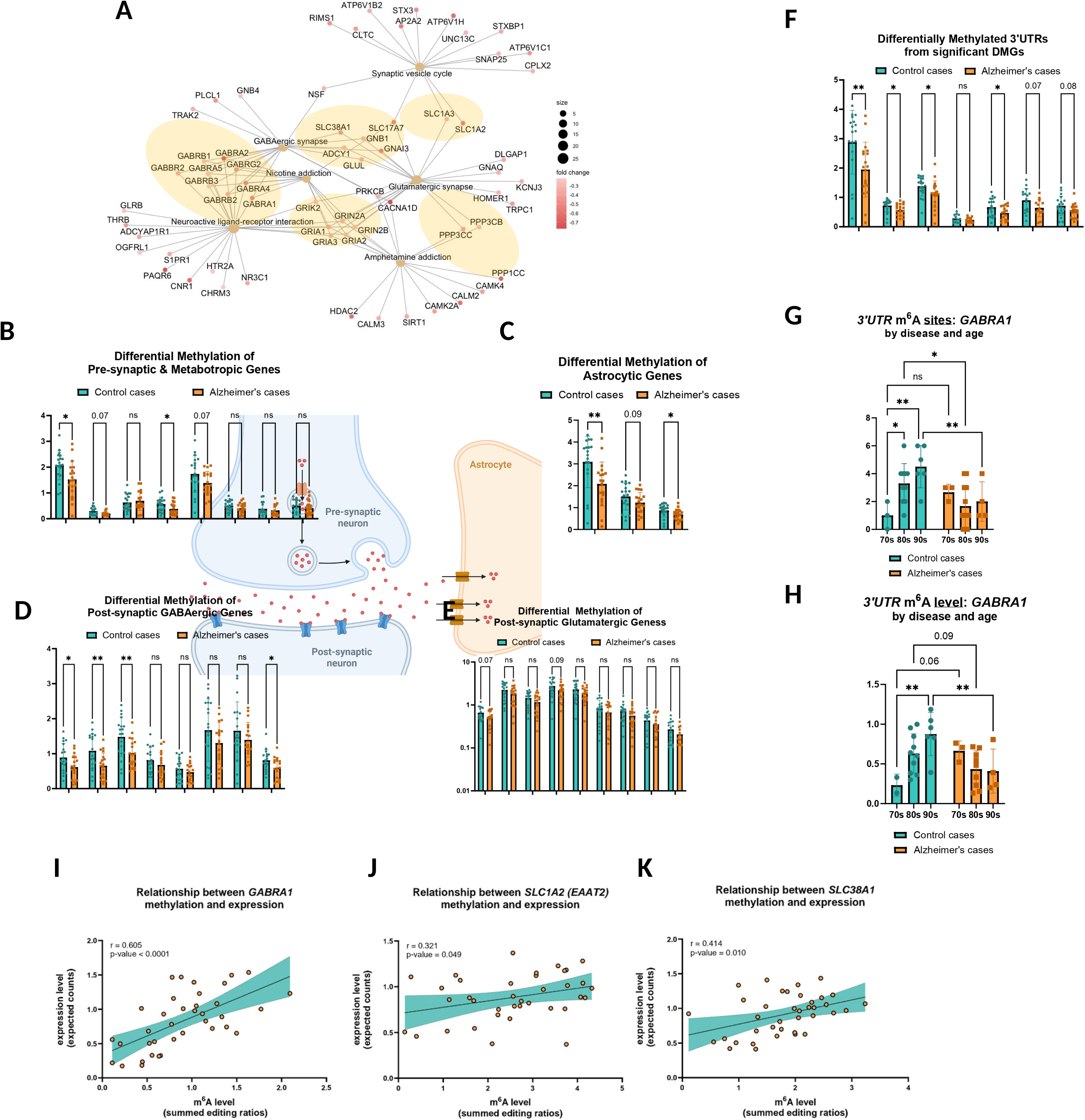
A tripartite synaptic network of differentially methylated genes (DMGs) is hypomethylated and downregulated in AD. (A) Network plot of DART-seq synaptic GSEA terms depicting shared genes between enrichment groups. Group size of each GSEA term and log2 fold change of each gene are shown by circle size and color gradient, respectively. (B) Comparison of m^6^A levels of pre-synaptic and metabotropic genes identified in the network plot of A between Control and AD cases. (* < 0.05) *P* values by unpaired Student’s *t* test. ns = not significant. Error bars represent mean ± standard deviation. (C) Comparison of m^6^A levels of astrocytic genes identified in the network plot of A between Control and AD cases. (* < 0.05, ** < 0.01) *P* values by Mann-Whitney U test. Error bars represent mean ± standard deviation. (D) Comparison of m^6^A levels of post-synaptic GABAergic genes identified in the network plot of A between Control and AD cases. (* < 0.05, ** < 0.01) *P* values by unpaired Student’s *t* test. ns = not significant. Error bars represent mean ± standard deviation. (E) Comparison of m^6^A levels of post-synaptic glutamatergic genes identified in the network plot of A between Control and AD cases. *P* values by unpaired Student’s *t* test. ns = not significant. Error bars represent mean ± standard deviation. (F) Comparison of 3’UTR-specific m^6^A levels of significant differentially methylated genes (DMGs) from B-E between Control and AD cases. (* < 0.05, ** < 0.01) *P* values by unpaired Student’s *t* test. ns = not significant. Error bars represent mean ± standard deviation. (G) Comparison of 3’UTR-localized m^6^A site quantification of *GABRA1* between Control and AD cases grouped by age. (* < 0.05, ** < 0.01) *P* values were generated by two-way ANOVA with Tukey’s HSD post-hoc test. ns = not significant. Error bars represent mean ± standard deviation. (H) Comparison of 3’UTR-specific m^6^A level quantification of *GABRA1* between Control and AD cases grouped by age. (* < 0.05, ** < 0.01) *P* values were generated by two-way ANOVA with Tukey’s HSD post-hoc test. ns = not significant. Error bars represent mean ± standard deviation. (I) Correlation of *GABRA1* m^6^A level (summed editing ratios from DART-seq) and expression level (expected counts from RNA-seq) using all brain samples. Regression is fitted to the equation: Y = 0.5447*X + 0.3386, has a positive Pearson correlation coefficient of 0.605, and is statistically significant (*P* < 0.0001). (J) Correlation of *SLC1A2 (EAAT2)* m^6^A level (summed editing ratios from DART-seq) and expression level (expected counts from RNA-seq) using all brain samples. Regression is fitted to the equation: Y = 0.06930*X + 0.7061, has a positive Pearson correlation coefficient of 0.321, and is statistically significant (*P* = 0.0492). (K) Correlation of *SLC38A1* m^6^A level (summed editing ratios from DART-seq) and expression level (expected counts from RNA-seq) using all brain samples. Regression is fitted to the equation: Y = 0.1747*X + 0.5979, has a positive Pearson correlation coefficient of 0.4144, and is statistically significant (*P* = 0.0098).

The strong age-associated differential increase in m^6^A sites (**Fig. 1G**) prompted us to examine this phenomenon among the tripartite synapse genes. First, we uncovered several genes in the pre-synapse, astrocyte, and post-synapse that are hypomethylated in AD (**Fig. 4B-E**). Then, we identified genes with differential 3’UTR-m^6^A level, given that we found the strongest age-related m^6^A effect among 3’UTR sites (**Fig. 1H**, **4F**). GABAergic receptors, notably *GABRA1*, and *SLC38A1* showed striking age-related increases in m^6^A methylation, but only in Control cases (**Fig. 4G-H, S11A-F**). In contrast, AD cases showed no change in m^6^A levels with age, recapitulating the aforementioned disease effect of m^6^A.

We proceeded to examine how the increase in m^6^A methylation impacts tripartite synapse gene expression. Expression of tripartite synapse genes is largely reduced in AD (**Fig. S12A, see Table S10**). Linear regressions among all cases demonstrated a strong link between m^6^A methylation and transcript expression for the GABAergic genes and glutamate/glutamine transporters *SLC1A2* and *SLC38A1*, but not for *GLUL* (**Fig. 4I-K, S12B-D**); increased m^6^A methylation correlated significantly with increased transcript expression. Non-synaptic genes, such as *MALAT1*, *HNRNPA2B1*, and *G3BP1*, showed no significant differential methylation between Control and AD cases (**Fig. S12E**) and did not show significant correlations between methylation and expression (**Fig. S12F-H**), suggesting that the relationship between differential methylation and expression is selective for genes at the tripartite synapse. The specific transcriptional patterns of m^6^A regulation are notable for occurring among transcripts that would mitigate increased glutamatergic signaling, potentially providing synaptic resilience in cognitively normal individuals. In contrast, deleterious glutamatergic signaling is a well-known feature of the pathophysiology of AD, which is potentially reflected by the absence of age-linked 3’UTR*-*m^6^A regulation in mitigating the reduction of GABAergic and glutamate-transporting genes among AD cases.^51,52^

### 3.8 Insufficient methylation of *GABRA1* in Alzheimer’s disease abolishes the association of m^6^A to GABRA1 fate

Our strongest differential, age-related change in m^6^A level was found on *GABRA1*, which was only one of twenty-two genes that exhibited transcriptome-wide significant decreases in expression and methylation in AD (**Fig. S12I, see Table S8 and Table S10**). To explore this relationship, we performed DART-PCR and Sanger sequencing of the 3’UTR m^6^A sites, given the strong differential relationship uncovered across aging (**Fig. 1H**), qPCR, and immunoblot experiments to link methylation, expression, and protein level with aging. There are seven 3’UTR m^6^A sites on *GABRA1* that were identified by our initial transcriptome-wide DART-seq approach, which showed variable levels of hypomethylation in AD by targeted DART-PCR and Sanger sequencing (**Fig. 5A, see Table S5**). These sites also had variable degrees of C>U editing that were specific to cytidines only in DRACH motifs (**Fig. 5B**). After summing the m^6^A levels of these sites, we confirmed the age-dependent increase of 3’UTR m^6^A level of *GABRA1* in Control cases, which was lost in AD cases (**Fig. 5C**). qPCR of total *GABRA1* level also matched our previous RNA-seq results showing 50% downregulation in AD cases compared to Control cases (**Fig. 5D**, **see Table S10**). However, we did not observe an effect of age at the transcript level here (**Fig. 5E**), suggesting that the age-effect between Control and AD cases is methylation-specific. We performed synaptosome isolation on our Control and AD cases (**Fig. 5F, Table S13**) and confirmed enrichment of GABRA1 protein in the synaptosomal fraction over the total homogenate fraction (**Fig. 5G**). We quantified a significant decrease in synaptosomal GABRA1 protein in AD cases compared to Control cases (**Fig. 5H**); no effect of age on synaptic GABRA1 protein level was observed (**Fig. 5I**), again establishing a methylation-specific effect of age between Control and AD cases.

**FIGURE 5:**
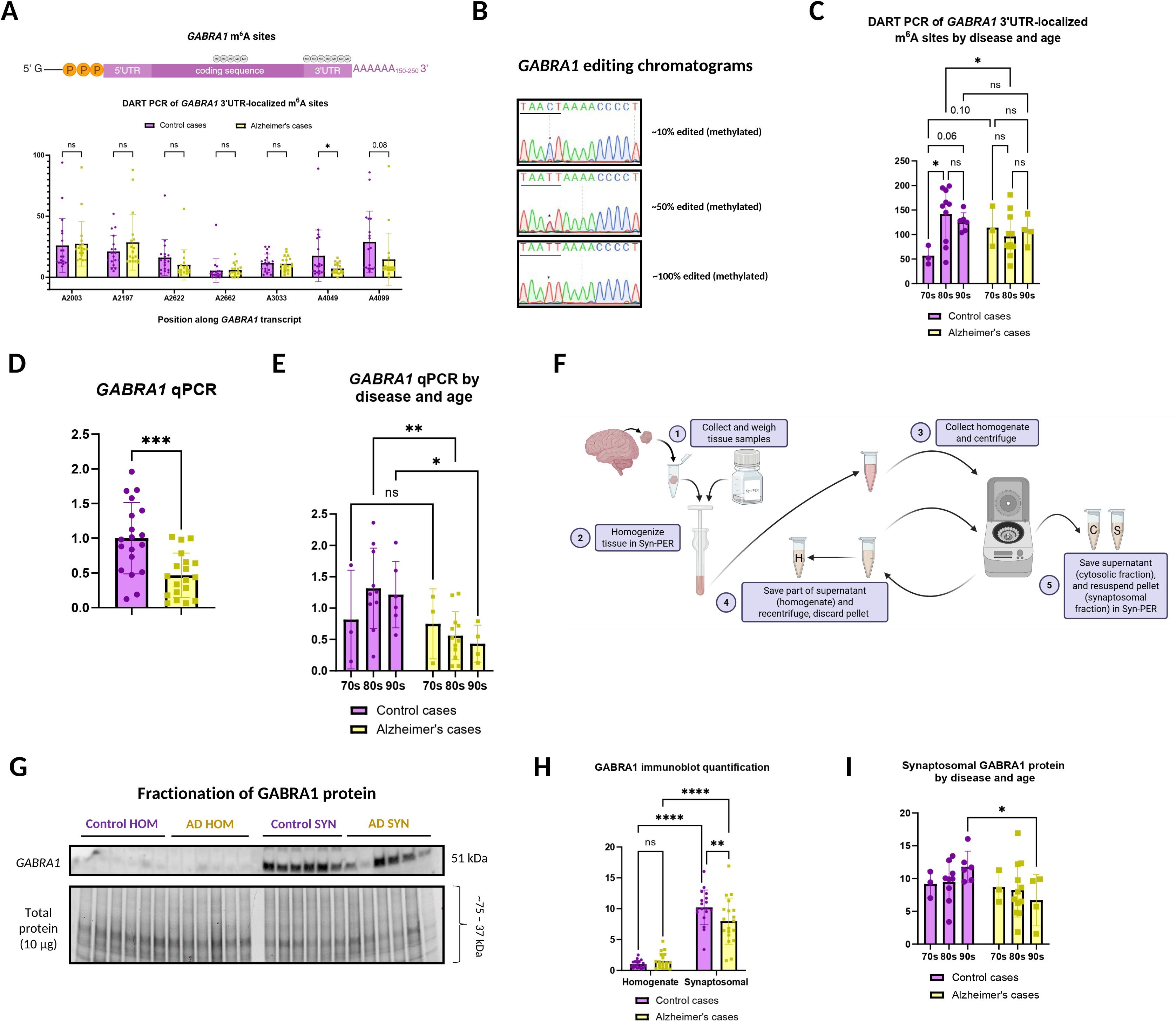
*GABRA1* m^6^A, expression, and protein levels all positively correlate and are reduced in AD. (A) Top: Diagram of *GABRA1* transcript depicting where m^6^A sites were identified, represented as gray “Me” circles. Total sites are listed in Table S5. Bottom: Comparison of m^6^A level across the 7 3’UTR*-*localized m^6^A sites of *GABRA1* by DART-PCR between Control and AD cases. * *P* < 0.05 for A4049 by unpaired Student’s *t* test. Error bars represent mean ± standard deviation. (B) Chromatograms showing various levels of *GABRA1* C>U editing. Underlined sequence represents identified DRACH motif. * denotes the C>U editing adjacent to the m^6^A-modified adenosine. Images were generated in SnapGene. (C) Comparison of 3’UTR-specific m^6^A level quantification in *GABRA1* between Control and AD cases grouped by age decades (70s, 80s, 90s). C>U editing ratio percentages from B were summed for each Control or AD case across the 7 3’UTR m^6^A sites. (* < 0.05) *P* values were generated by two-way ANOVA with Tukey’s HSD post-hoc test. Error bars represent mean ± standard deviation. (D) Comparison of *GABRA1* expression normalized to *GAPDH* expression by qPCR using the 2^-ΔΔ*CT*^ method between Control and AD cases. *** *P* < 0.001 by unpaired Student’s *t* test. Error bars represent mean ± standard deviation. (E) Comparison of *GABRA1* expression between Control and AD cases (quantified in D) grouped by age. (* < 0.05, ** < 0.01) *P* values were generated by two-way ANOVA with Tukey’s HSD post-hoc test. Error bars represent mean ± standard deviation. (F) Diagram showing workflow of synaptosome extraction using Syn-PER. Tissue samples were weighed (1) and homogenized in Syn-PER (2). Homogenate was collected (3) and centrifuged twice: once to spin out insoluble debris and collect the homogenate (4) and the other to separate the cytosolic fraction (supernatant) from synaptosomal fraction (pellet, 5). (G) Immunoblot comparing GABRA1 protein level (top) between homogenate (HOM) and synaptosomal (SYN) fractions between Control and AD cases. Total protein scan (below) was used to normalize for protein input. Uncropped and synaptosomal validation immunoblots can be found in Table S12. (H) Quantification of homogenate and synaptosomal GABRA1 protein levels and comparison between Control and AD cases. (** < 0.01, **** < 0.0001) *P* values were generated by two-way ANOVA with Fisher’s LSD post-hoc test. Error bars represent mean ± standard deviation. (I) Comparison of GABRA1 synaptosomal protein level between Control and AD cases (quantified in H) grouped by age. (* < 0.05) *P* values were generated by two-way ANOVA with Fisher’s LSD post-hoc test. Error bars represent mean ± standard deviation.

We then generated correlation matrices to understand how the relationship between GABRA1 metabolism and age may be different for Control cases compared to AD cases. In Control cases, the 3’UTR m^6^A level of *GABRA1* positively associated with age, and m^6^A levels of various 3’UTR DRACH sites associated with *GABRA1* expression, protein level in the homogenate, and protein level in the synaptosome (**Fig. 6A**). Conversely in AD cases, total 3’UTR m^6^A level of *GABRA1* had no association with age, although individual 3’UTR DRACH sites did show an age-related correlation (**Fig. 6B**). Remarkably, the relationship of individually methylated *GABRA1* 3’UTR DRACH sites with expression or protein level of GABRA1 was also abolished. These associations suggest that a critical level of m^6^A may exist to elicit control over transcript metabolism, which in AD, could be too low to be met for *GABRA1* (**Fig. 6C**).

**FIGURE 6:**
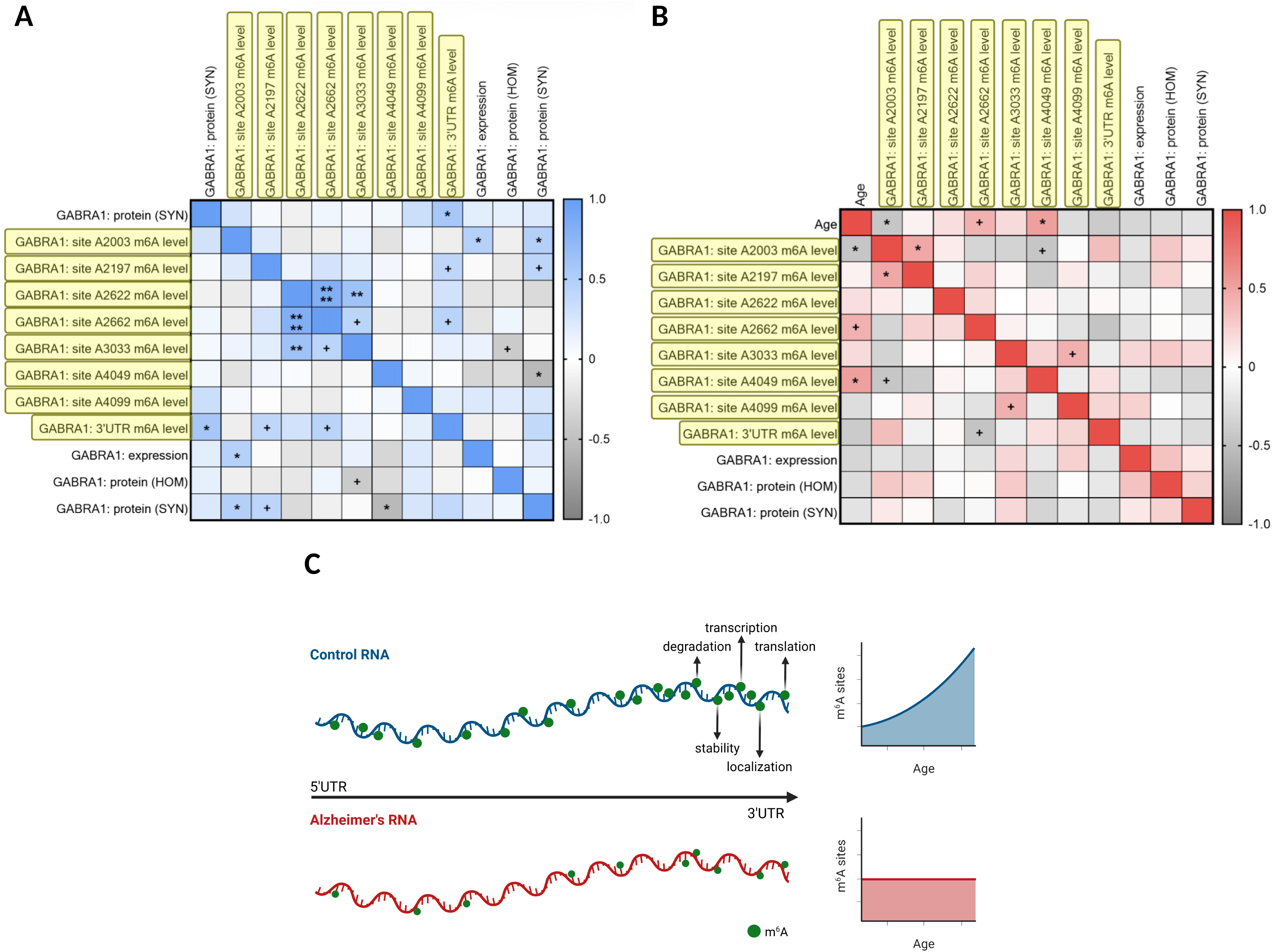
The association of site-specific *GABRA1* methylation to expression and protein levels is lost in Alzheimer’s disease. (A) Correlation matrix of age, m^6^A level (editing ratio) from DART-PCR of DRACH sites in the 3’UTR of *GABRA1*, *GABRA1* expression from qPCR, and GABRA1 protein levels from immunoblot quantification of Control cases. Pearson correlation coefficients range from -1.0 to 1.0. ^+^ *P* < 0.10, * *P* < 0.05, ** *P* < 0.01, **** *P* < 0.0001. HOM = homogenate fraction. SYN = synaptosomal fraction (B) Correlation matrix of age, m^6^A level (editing ratio) from DART-PCR of DRACH sites in the 3’UTR of *GABRA1*, *GABRA1* expression from qPCR, and GABRA1 protein levels from immunoblot quantification of AD cases. Pearson correlation coefficients range from -1.0 to 1.0. ^+^ *P* < 0.10, * *P* < 0.05. HOM = homogenate fraction. SYN = synaptosomal fraction. (C) Overview of m^6^A modification comparison between Control and AD transcripts. In Control cases, there are more m^6^A sites and higher levels of methylation, especially enriched in the 3’UTR, which positively associates with age. This level of methylation is sufficient to produce an association of m^6^A to transcript metabolism at the site-specific level. In AD cases, there are fewer m^6^A sites and lower levels of methylation, especially depleted in the 3’UTR, and do not associate with age. This level of methylation is insufficient and blocks the association of m^6^A to transcript metabolism at the site-specific level.

## 4 DISCUSSION

RNA modifications, such as m^6^A methylation, represent a key level of control for RNA metabolism. We present the first site-specific analysis of m^6^A changes in the human brain and characterize m^6^A changes in AD at the site, domain, and gene levels. We uncovered over 36,500 sites across 6,800 genes that were enriched for transcripts regulating synaptic activity (**Fig. 1D**). Brain samples from cognitively normal individuals exhibit increase in m^6^A sites with age, which was seen most strongly in 3’UTR-localized m^6^A sites (**Fig. 1H**). In contrast, AD brains show a stable number of m^6^A sites with aging and reduced levels of m^6^A-modifications in transcripts (**Figs. 1E, 2A**). For instance, *MAPT* exhibits hypomethylation in AD cases (**Fig. 2A**). The increase in m^6^A labeling with age is particularly apparent in the GABAergic/glutamatergic tripartite synapse, where m^6^A labeling correlates with expression level for GABA-A receptor α subunits *GABRA1*, *GABRA2*, and *GABRA*4, glutamate transporter *SLC1A2*. and glutamine transporter SLC38A1 (**Fig. 4I-K, S12B-C**). The absence of age-related adaptation of m^6^A methylation could indicate that AD pathophysiology disrupts the regulation of RNA metabolism through impaired m^6^A methylation and needs to be further investigated.

Aberrant m^6^A methylation in AD appears to associate with dysregulation of the tripartite synapse axis. Disease-linked changes in m^6^A sites occur on *GRIN2A*, *GRIN2B*, *GABRA1*, *GABRA2*, and *GABRA4* transcripts (**Fig. S8D**). This could indicate a disruption in neurotransmission by differential methylation of excitatory and inhibitory receptors at the site level. At the transcript level, *GABRA1*, *GABRA2*, *GABRA4*, *SLC1A2*, and *SLC38A1* all showed strong disease-linked hypomethylation in AD (**Fig. 4B-D**) and robust positive correlation between m^6^A labeling and transcript levels (**Fig. 4I-K, S12B-C**). Correlation was also apparent between GABRA1 protein and m^6^A level at particular *GABRA1* m^6^A sites (**Fig. 6A**), which suggests that m^6^A labeling, transcription, and translation of the tripartite synaptic genes in AD are interconnected. Reduction in methylation of tripartite GABA receptors and glutamate transporters might contribute to the detrimental excitatory signaling that occurs in AD models and AD brains.^53,54^ In contrast, cognitively normal Control brains exhibit progressive age-linked increases (across 70-, 80-, and 90-year-old cases) in m^6^A levels in the 3’UTR, as seen for *GABRA1*, *GABRA2*, *GABRA4*, *SLC1A2*, and *SLC38A1* transcripts (**Figs. 4G-H, S11A-C, S11E**). This implicates m^6^A in regulating GABAergic and glutamatergic signaling (**Figs. 1I, S5A**, **3A**, **4A**). Other studies noted increases in m^6^A with aging but have not previously identified the role of the tripartite synaptic genes with aging.^55,56^ Increased excitotoxic activity tends to occur with aging and disease.^57^ The age-related increase in m^6^A labeling of GABA receptors and glutamate transporters could reflect a protective compensatory response, which dampens excitatory activity and is dysregulated in AD.

The general dysregulation of m^6^A sites in the 3’UTR among AD cases suggests a widespread impairment of RNA metabolism, although it is important to note the CDS also contains many m^6^A sites (**Fig. 1B**). In the brain, m^6^A has been shown to influence transcript localization to synapses.^49,58^ The 3’UTR contains binding motifs that m^6^A readers and RBPs can use to direct transcript localization to and translation at synapses.^59^ In AD, 3’UTR-directed hypomethylation of transcripts encoding tripartite synapse genes (**Fig. 4F**) could indicate a reduction in synaptic trafficking of transcripts, reducing the pool of available transcripts necessary for activity-dependent translation, neuroplasticity, and protection from excitotoxicity. These endpoints should be explored in future studies.

Tauopathies, including AD, elicit dysfunction of proteostasis with a concomitant stress response, which was reflected in the m^6^A-transcriptome. Methylation of multiple related stress response transcripts was dysregulated in AD, including the translation regulators *EIF2AK2*, *EIF1AX*, *EIF4B*, and *EIF4BP3* (**Figs. 2A, S9D, S9H, S8D**). The DAS for *EIF2AK2* shows more consistent hypermethylation among AD cases than Control cases (**Fig. S8D**). This is striking given that more sites and genes are hypomethylated than hypermethylated in AD (**Fig. S8C, 2A**), suggesting an increase in stress-dependent regulation of translation in AD. Previous studies support the idea that m^6^A is highly responsive to neuronal conditions; for instance, many studies show that RNA translation decreases with stress and disease, and activity-dependent translation of m^6^A-modified transcripts is also well-documented in the brain.^3,18,49^ We also observed DAS for multiple *HSP90* paralogs, which might reflect impairment of proteostasis in AD with a resulting exacerbation of pathologies. Genes that regulate ubiquitin-mediated proteolysis, including ubiquitin conjugating enzyme genes (*UBE2D3*, *UBE2K*, *UBE2G1*) and Cullin family genes (*CUL4B*, *CUL5*, *CUL2*) are depleted in both the DART-seq and RNA-seq analyses (**Figs. 3A-C**, **3G**). Thus, m^6^A-linked downregulation of these genes might be associated with dysfunction of ubiquitin signaling and proteostasis, reducing protein clearance and potentially enhancing protein aggregation.

Many genes associated with AD pathophysiology showed differential m^6^A levels. For example, *MAPT* is hypomethylated (**Fig. 2A**) and the m^6^A level at DRACH site A5942 of *MAPT* negatively correlates with MAPT protein level. This suggests that in AD, reduction of m^6^A on *MAPT* is linked with increased tau protein. Genes such as *FYN* (a kinase that phosphorylates tau at synapses) and *PPP1CB* (part of a tau phosphatase complex) have DAS in AD cases (**Fig. S8D**), linking m^6^A to tau phosphorylation status.^60,61^ Interestingly, the DAS on *FYN* (chr6:111780572, **Table S7**) is a common SNP (rs12910, 0.47; NCBI dbSNP) that is associated with increased risk of AD in the Chinese Han population.^62^ Other genes that are associated with AD pathophysiology and showed differential m^6^A sites include the proteostasis genes *HSP90AA1, HSP90B1, HSP90AB1, HIF1A*, and the translational stress response genes *EIF2AK2*, *G3BP1*, *DDX5*, *HNRNPC* and *CAPRIN1; G3BP1* and *CAPRIN1* are essential for formation of stress granules.^3^ These findings link aberrant m^6^A methylation at the site and gene levels to well established players involved in AD pathophysiology. Restoring methylation of these sites to non-AD levels could offer a mechanism of AD intervention.

The analyses reported in this manuscript provide a benchmark for understanding how m^6^A methylation responds to aging and neurodegeneration in humans. The nucleotide level analysis for the human m^6^A epitranscriptome provides an important reference for interpreting other studies using less precise approaches, such as MeRIP-seq and/or studies that use animal models that incompletely reflect AD pathophysiology. The reduced precision of MeRIP-seq combined with differences between the human condition and animal models might contribute to variable results among m^6^A studies in the literature. A small study of human brain samples (n=6, using MeRIP-seq) shows similar results with increased m^6^A methylation of mRNA in the normal aged human brain region Brodmann Area 10 (comparable to the brain region used for this study).^18^ Studies of aging in Drosophila also indicate that m^6^A increases with age in the brain.^63^ Mixed results of m^6^A in aging were observed among studies using mouse models of AD that express human APP/PS1 or human APP (mut) knock-in/P301S and Drosophila models expressing human R406W tau.^64–66^ Total m^6^A appears to increase, but m^6^A methylation of specific mRNA transcripts tends to be lower. Our prior work also demonstrated an m^6^A increase with disease severity among neurons showing AD pathology.^8^ However, these results relied on antibody-based imaging and ELISA experiments, which do not discriminate between RNA species. Given the high relative abundance of rRNA to mRNA, it is possible that the general increase in m^6^A labeling observed with m^6^A-targeting antibodies reflects increased m^6^A methylation of rRNA.^8^ The observations from this study provide the necessary granular detail for the human brain mRNA transcriptome that identifies a distinct, robust age-related regulation of m^6^A in normal Control subjects that contrasts with dysregulation in AD subjects.

The current study has inherent limitations to consider. First, DART constructs are limited by the YTH binding capacity and restricted to m^6^A site identification within DRACH motifs. To circumvent false positives and address binding capacity limitations, we used the APOBEC1-YTH^mut^ construct as a negative control benchmark. Although most m^6^A sites on mRNA occur within the DRACH context, non-DRACH m^6^A sites exist, which can be detected by other methods such as GLORI-seq.^67,68^ If scDART methods are developed for use with fixed tissue, studies of cellular heterogeneity become possible; such methods are currently available for use *in vitro* and *in vivo*.^27,32^ Second, we did not observe a strong m^6^A site enrichment around the stop codon (**Fig. 1C**).^69^ However, our m^6^A site distribution is similar to what prior DART-seq studies reported.^15,48^ Differences in m^6^A sequencing technique, species (e.g. human versus mouse), and sample quality (e.g., cell lysate versus flash frozen tissue) could impact the m^6^A site distribution detected.^11^

The results presented in this study demonstrate a striking response of RNA metabolism to the pathophysiology of AD, and delineate how RNA modifications, especially m^6^A, might alter RNA metabolism in the brain with respect to age and disease. We uncovered a strong 3’UTR m^6^A component in normal aging, a global reduction of m^6^A-modified transcripts in AD, and two m^6^A-associated dysregulated axes at tripartite synapses and proteostasis. The results provide granularity to m^6^A methylation that was previously unknown, such as with site-specific differential changes in AD. Knowing the specific m^6^A methylation changes occurring with AD could lead to future approaches that clarify disease heterogeneity as well as engineered approaches to reverse m^6^A modifications in specific gene targets contributing to the pathophysiology of AD.

## AUTHOR CONTRIBUTIONS

JLL conceptualized the study, supported bioinformatics analyses, completed and oversaw molecular experiments, wrote the original draft, and co-wrote the revised draft. JTMA completed revision experiments, supported bioinformatics analyses, and co-wrote the revised draft. JH and TN led bioinformatics analyses, with TN supporting molecular experiments. ZW supported bioinformatics analyses and figure generation. SFJS provided materials and reagents for molecular experiments. KS, LD, KT, YM, and JGR supported molecular experiments. JM supported bioinformatics analyses. SP and AK provided conceptual support. NMK, PTN, KM, and JTCW provided materials and reagents for molecular experiments. LH supported bioinformatics analyses. XZ oversaw bioinformatics analyses. BW supported study conceptualization, data interpretation and reviewed and edited all drafts.

## ACKNOWLEDGEMENTS

The sequencing was performed at the Boston University Medical Campus Microarray and Sequencing Resource.

## DECLARATION OF GENERATIVE AI AND AI-ASSISTED TECHNOLOGIES IN THE WRITING PROCESS

During the preparation of this work, JLL used ChatGPT in order to assist with writing the Abstract and adhere to word count restrictions. After using this service, JLL reviewed and edited the content as needed and takes full responsibility for the content of the publication.

## DATA AND MATERIALS AVAILABILITY

The DART-seq data generated in this study have been deposited in the NCBI Gene Expression Omnibus (GEO) under accession number GSE297551. Source data are provided with this manuscript. All scripts used to generate data for this study are available in the GitHub repository: https://github.com/WolozinLab/Dart-seq_data_processing_code.

## CONSENT STATEMENT

Consent was not necessary for the studies presented in this manuscript.

---

**Supplemental Figure 1:**
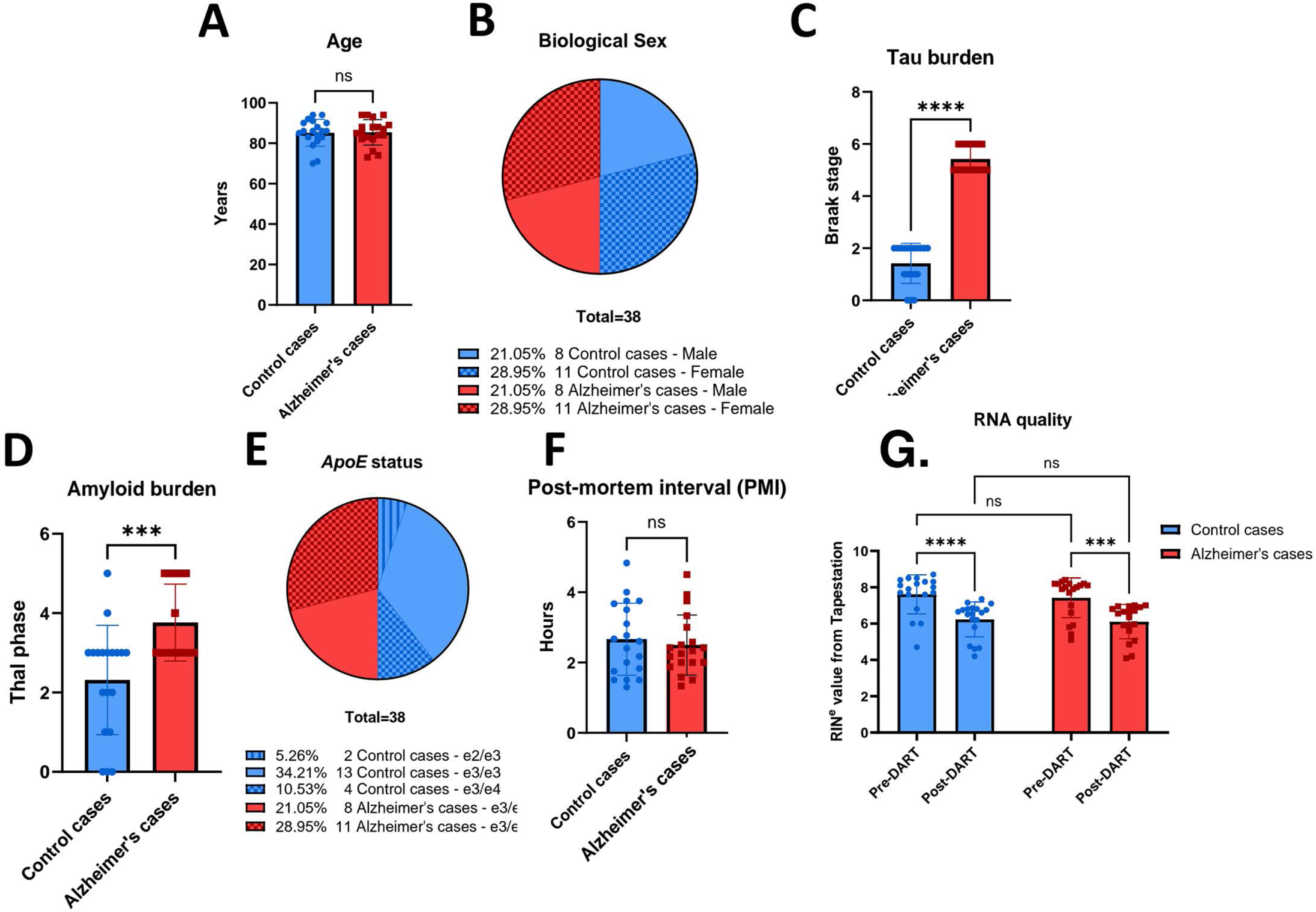
Post-mortem human brain samples are of high quality and exhibit distinct tau and amyloid pathologies between Control and AD cases. (A) Comparison of age (years) between Control and AD cohorts. *P*>0.05 by unpaired Student’s t test. Error bars represent mean± standard deviation. (B) Pie chart of biological sex between Control and AD cohorts. (C) Comparison of tau burden (Braak stage) between Control and AD cohorts. **** *P<* 0.0001 by Mann-WTutney U test. Enor bars represent mean = standard deviation. (D) Comparison of amyloid bur den (Thal phase) between Control and AD cohorts. *** *P<* 0.001 by Mann-Whitney U test. Error bars represent mean± standard deviation. (E) Pie chart of *APOE* status between Control and AD cohorts. (F) Comparison of post-mortem interval (hours) between Control and AD cohorts. P>0.05 by unpaired Student’s r test. Error bars represent mean÷ standard deviation. (G) Comparison of RNA quality (RIN°) between Contr ol and AD cohorts before and after DART reactions. **** *P* < 0.0001 by two-way ANOVA with Fisher’s LSD post-hoc test. Error bars represent mean≈ standard deviation.

**Supplemental Figure 2:**
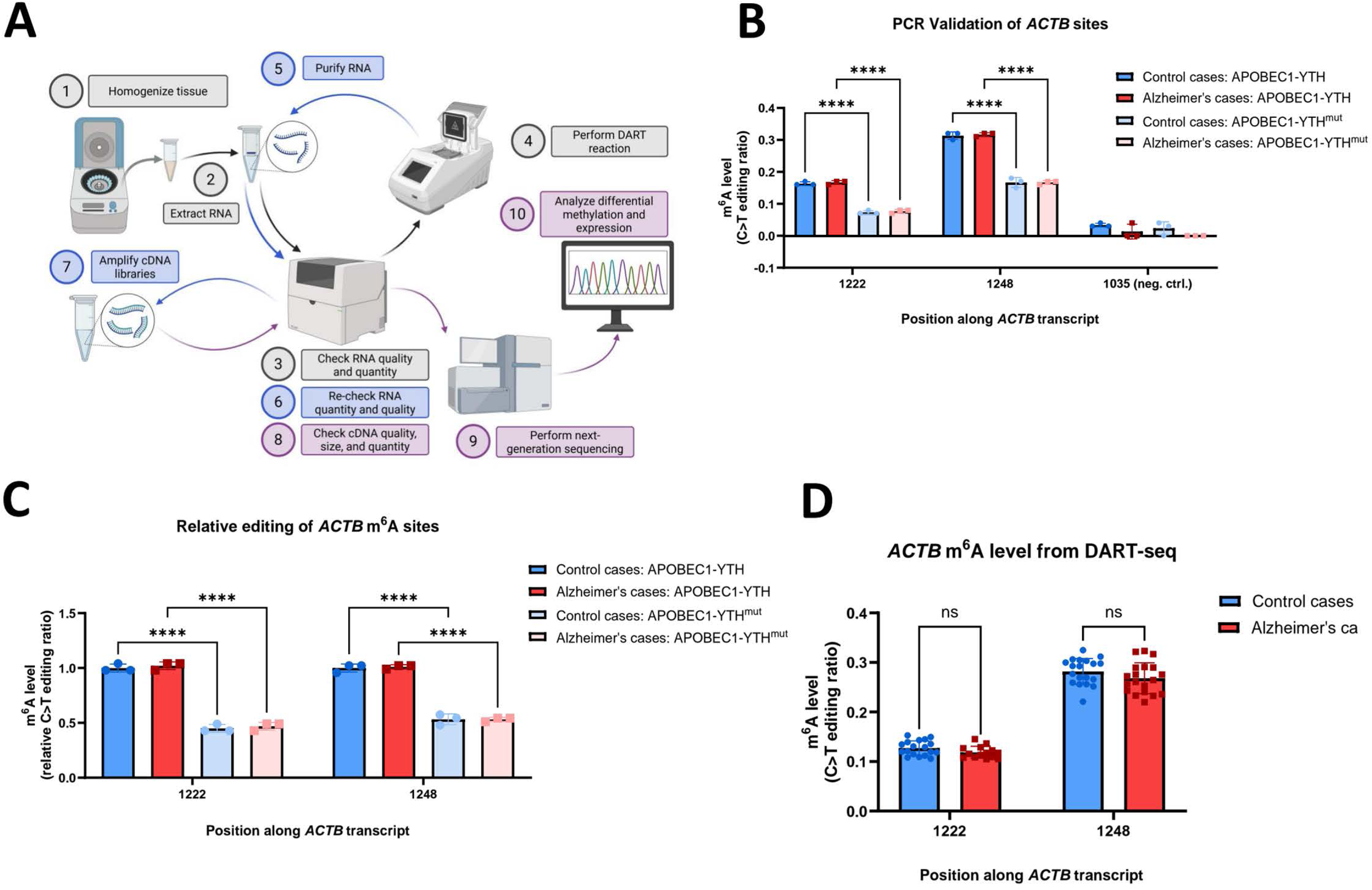
Overview of DART workflow and validation of ACTB m^6^A sites. (A) Diagram of DART workflow steps: 1) Homogenization, 2) RNA extraction, 3) Tapestation check, 4) DART reaction, 5) RNA repurification, 6) Tapestation check, 7) cDNA library construction (or DART-PCR), 8) Tapestation check, 9) NGS (or Sanger sequencing), 10) Differential expression and methylation analysis. (B) Quantification of m^6^A level (OU editing ratio) from DART-PCR and Sanger sequencing of *ACTB* m^6^A sites with negative control between Control and AD cohorts treated with either APOBEC l-YTH or AOBEC1 -YTH““*. **** *P* < 0.0001 by two-way ANOVA with Tukey’s HSD post-hoc test for site 1222 and site 1248. *P* > 0.05 by two-ANOVA with Tukey’s HSD post-hoc test for site 1035 (negative control). Error bars represent mean± standard deviation. (C) Quantification of relative m^6^A level (normalized OU editing ratio) from DART-PCR and Sanger sequencing of *A CTB* m^6^A sites between Control and AD cohorts treated with either APOBEC1-YTH or APOBECl-YTH““‘. **** *P* < 0.0001 by two-way ANOVA with Tukey’s HSD post-hoc test for site 1222 and site 1248. Error bars represent mean ± standard deviation. (D) Comparison of m^6^A level (OU editing ratio) of *ACTB* m^6^A sites between Control and AD cohorts from DART-seq. P>0.05 by unpaired Student’s *t* test for both sites. Error bars represent mean± standard deviation.

**Supplemental Figure 3:**
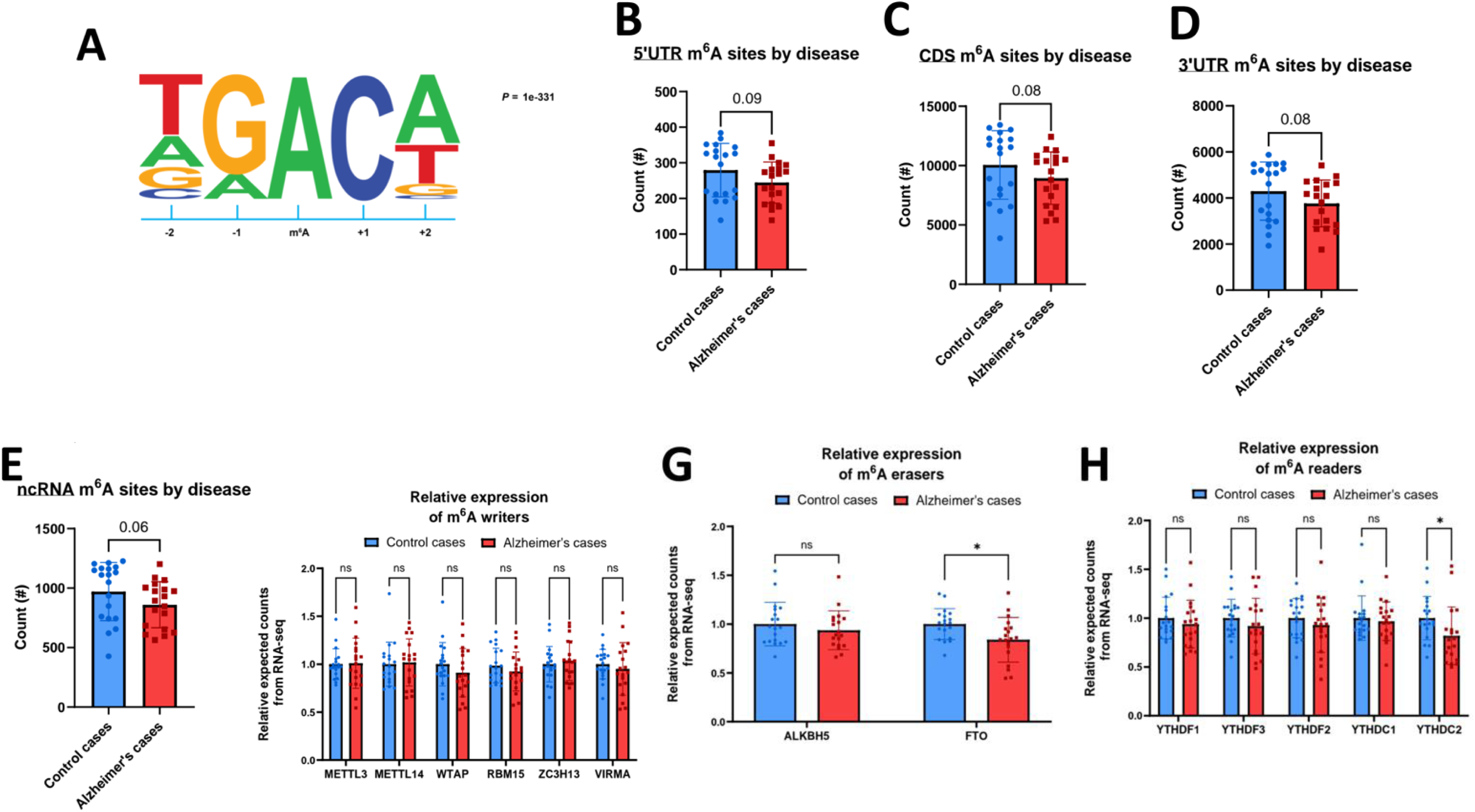
Profiling of total m^6^A sites. (A) DRACH motif enrichment of m^6^A sites. (B) Comparison of 5’UTR-localized m^6^A site quantification between Control and AD cases. *P* = 0.0891 by Mann-Whitney U test. Error bars represent mean ± standard deviation. (C) Comparison of CDS-localized m^6^A site quantification between Control and AD cases. *P* = 0.0797 by Mann-Whitney U test. Error bars represent mean ± standard deviation. (D) Comparison of 3’UTR-localized m^6^A site quantification between Control and AD cases. *P* = 0.0797 by Mann-Whitney U test. Error bars represent mean ± standard deviation. (E) Comparison of ncRNA-specific m^6^A site quantification between Control and AD cases. *P* = 0.0601 by Mann-Whitney U test. Error bars represent mean ± standard deviation. (F) Comparison of expression level for m^6^A writers between Control and AD cases. *P* values were generated by unpaired Student’s t test. Error bars represent mean ± standard deviation. (ns = not significant). (G) Comparison of expression level for m^6^A erasers between Control and AD cases. *P* values were generated by unpaired Student’s t test. Error bars represent mean ± standard deviation. (* < 0.05, ns = not significant). (H) Comparison of expression level for m^6^A readers between Control and AD cases. *P* values were generated by unpaired Student’s t test. Error bars represent mean ± standard deviation. (* < 0.05, ns = not significant).

**Supplemental Figure 4:**
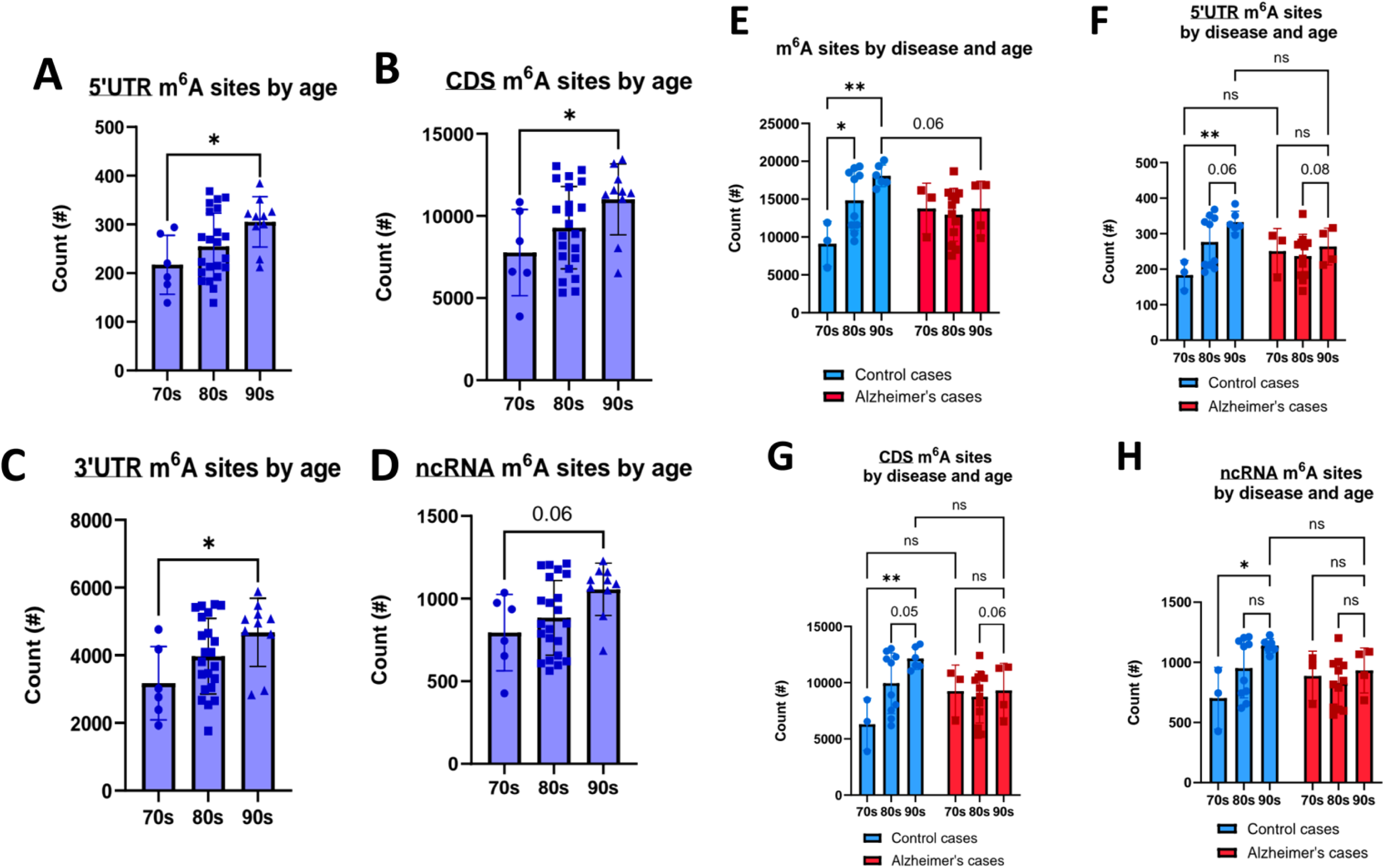
Profiling of domain-specific m^6^A sites by disease only, age only, and by disease and age combined. (A) Comparison of 5’UTR-localized m^6^A site quantification of all cases grouped by age decade (70s, 80s, and 90s). * *P* < 0.05 by one-way ANOVA with Tukey’s HSD post-hoc test. Error bars represent mean ± standard deviation. (B) Comparison of CDS-localized m^6^A site quantification of all cases grouped by age. * *P* < 0.05 by one-way ANOVA with Tukey’s HSD post-hoc test. Error bars represent mean ± standard deviation. (C) Comparison of 3’UTR-localized m^6^A site quantification of all cases grouped by age. * *P* < 0.05 by one-way ANOVA with Tukey’s HSD post-hoc test. Error bars represent mean ± standard deviation. (D) Comparison of ncRNA-specific m^6^A site quantification of all cases grouped by age. *P* = 0.0560 by one-way ANOVA with Tukey’s HSD post-hoc test. Error bars represent mean ± standard deviation. (E) Comparison of m^6^A site quantification between Control and AD cases grouped by age. (* < 0.05, ** < 0.01) *P* values were generated by two-way ANOVA with Tukey’s HSD post-hoc test. Error bars represent mean ± standard deviation. (F) Comparison of 5’UTR-localized m^6^A site quantification between Control and AD cases grouped by age. (** < 0.01) *P* values were generated by two-way ANOVA with Tukey’s HSD post-hoc test. Error bars represent mean ± standard deviation. (G) Comparison of CDS-localized m^6^A site quantification between Control and AD cases grouped by age. (** < 0.01) *P* values were generated by two-way ANOVA with Tukey’s HSD post-hoc test. Error bars represent mean ± standard deviation. (H) Comparison of ncRNA-specific m^6^A site quantification between Control and AD cases grouped by age. (* < 0.05) *P* values were generated by two-way ANOVA with Tukey’s HSD post-hoc test. Error bars represent mean ± standard deviation.

**Supplemental Figure 5:**
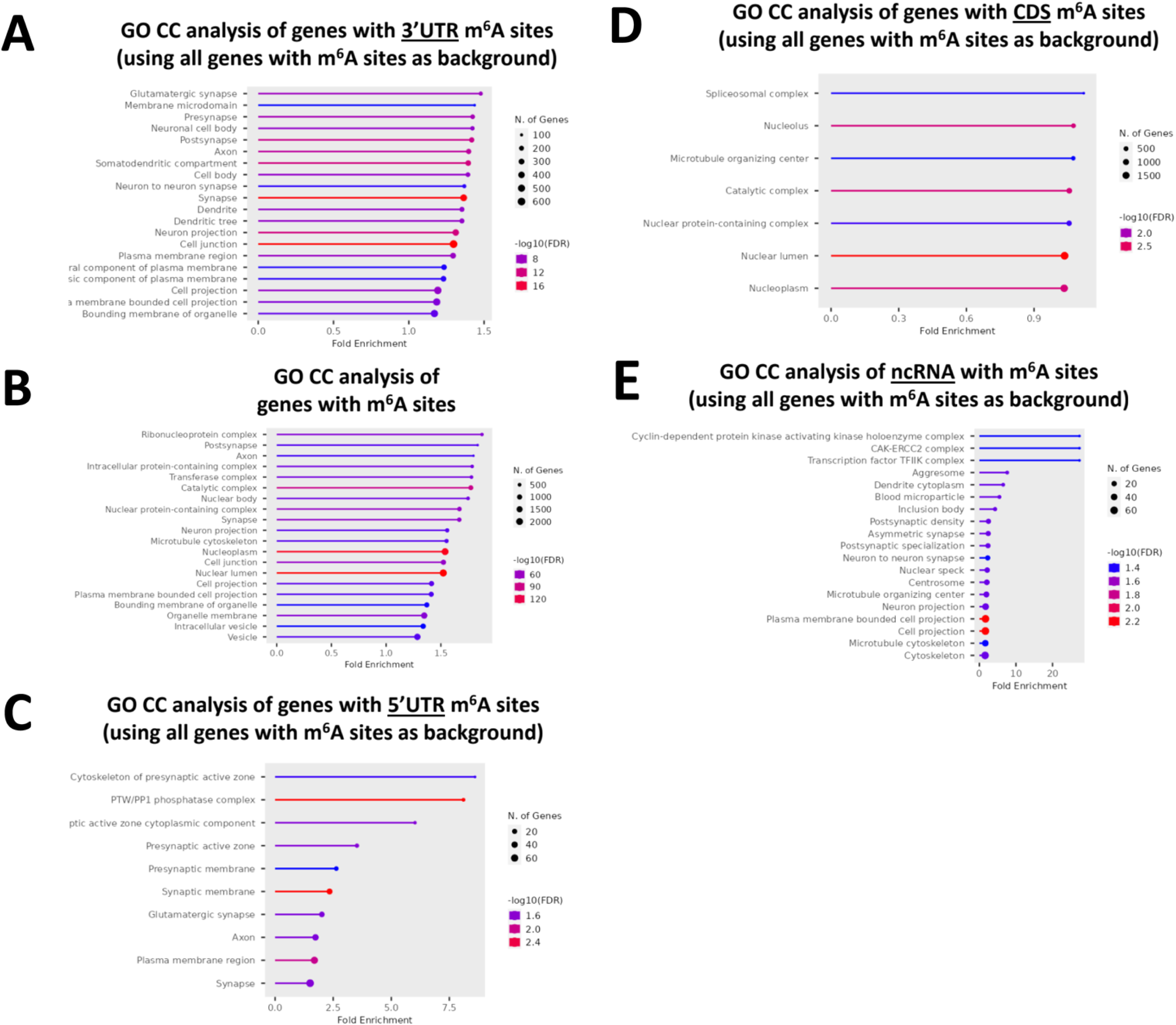
Gene ontology (GO) cellular compartment (CC) analyses reveal strong synaptic enrichment of genes with 3’UTR m^6^A sites. (A) GO CC analysis of genes with 3’UTR-localized m^6^A sites. All genes with m^6^A sites, regardless of site location, were used as a background reference. (B) GO CC analysis of genes with m^6^A sites. (C) GO CC analysis of genes with 5’UTR-localized m^6^A sites. All genes with m^6^A sites, regardless of site location, were used as a background reference. (D) GO CC analysis of genes with CDS-localized m^6^A sites. All genes with m^6^A sites, regardless of site location, were used as a background reference. (E) GO CC analysis of genes with ncRNA-specific m^6^A sites. All genes with m^6^A sites, regardless of site location, were used as a background reference.

**Supplemental Figure 6:**
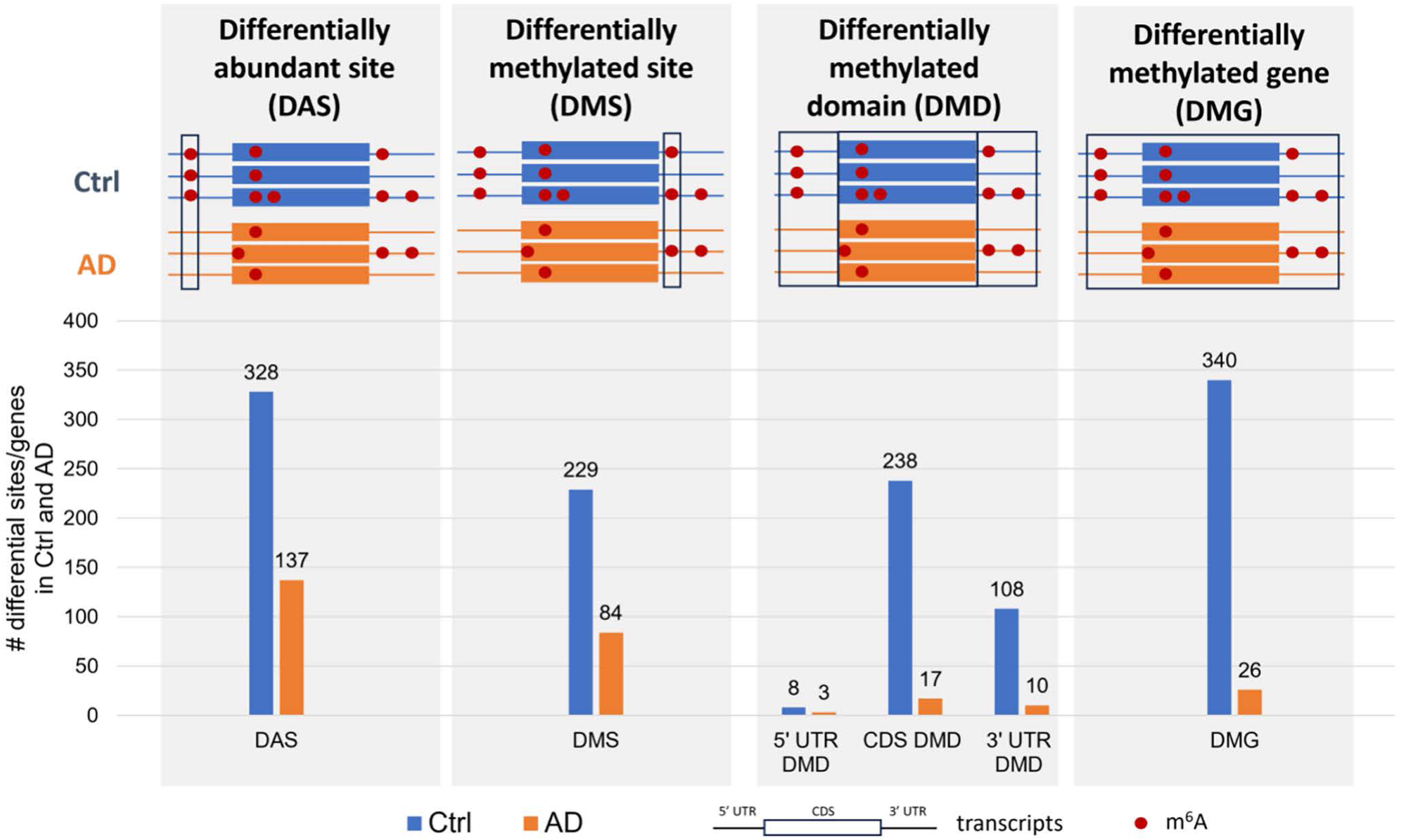
Quantification for differential methylation using different parameters between AD and Control. Bar chart showing numbers for differentially abundant sites (DAS, **Table S7**). differentially methylated sites (DMS. **Table S6**). differentially methylated domains (DMD) and differentially methylated genes (DMG, **Table S8**). Sites were differentially abundant in Control cases if >50% of cases had a site and <10% of AD cases had a site. Conversely. DAS in AD cases had >50% representation and <10% representation in Control cases. Methylation level of all sites, regardless of case representation, were summed and averaged across an entire gene or gene domain (5’UTR. CDS, 3’UTR) and subjected to differential analysis.

**Supplemental Figure 7:**
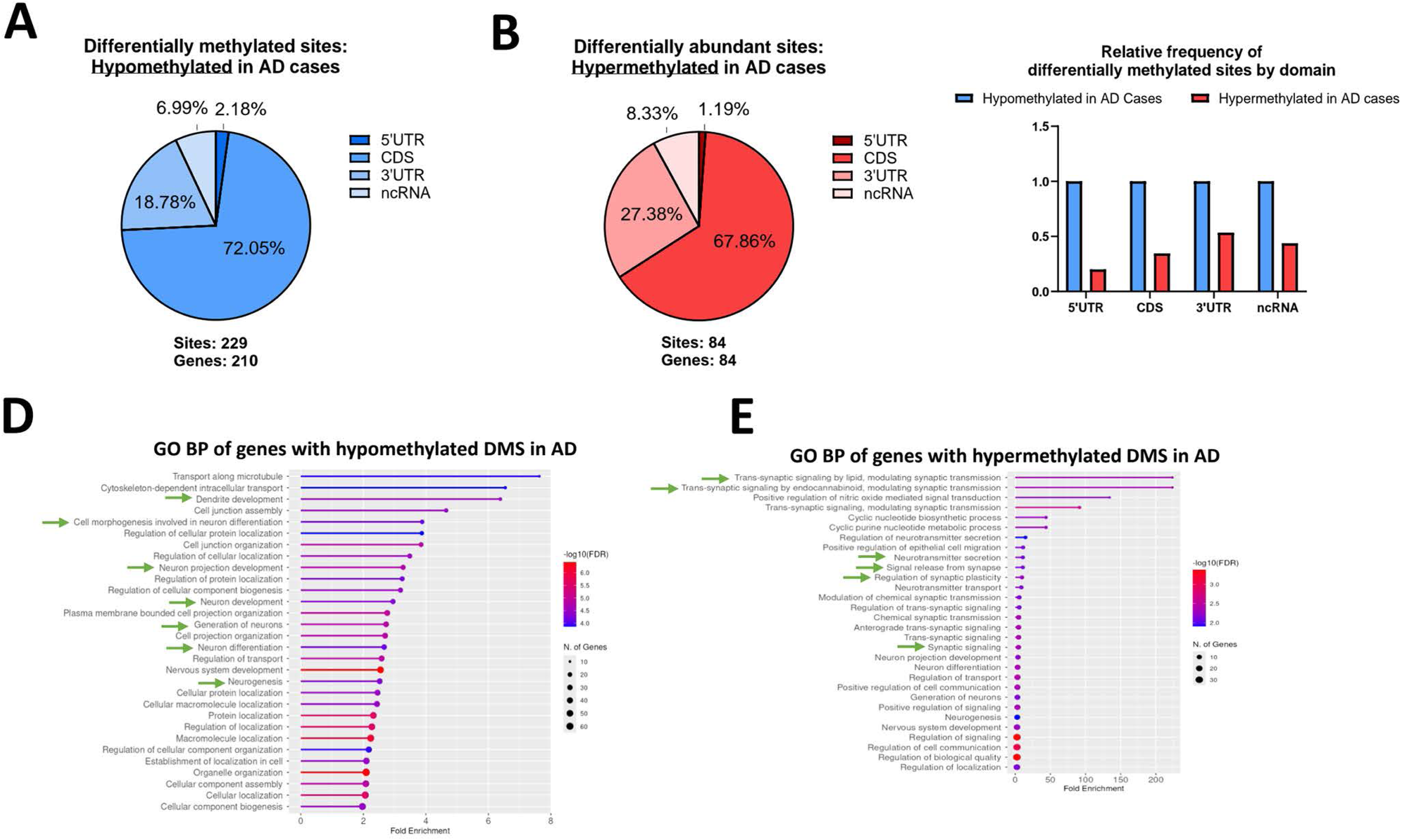
Differentially abundant site (DAS) characterization by domain distribution and gene ontology enrichment. (A) Pie chart showing percentage of hypomethylated m^6^A sites along different mRNA domains and on ncRNA. (B) Pie chart showing percentage of hypermethylated m^6^A sites along different mRNA domains and on ncRNA. (C) Bar plot showing distribution of hypo-and hypermethylated sites by mRNA domains and on ncRNA. (D) GO Biological Process analysis of genes with AD hypomethylated DMS sites. Terms related to neuron development and synaptic function are marked by green arrows. (E) GO Biological Process analysis of genes with AD hypermethylated DMS sites. Terms related to neuron development and synaptic function are marked by green arrows.

**Supplemental Figure 8:**
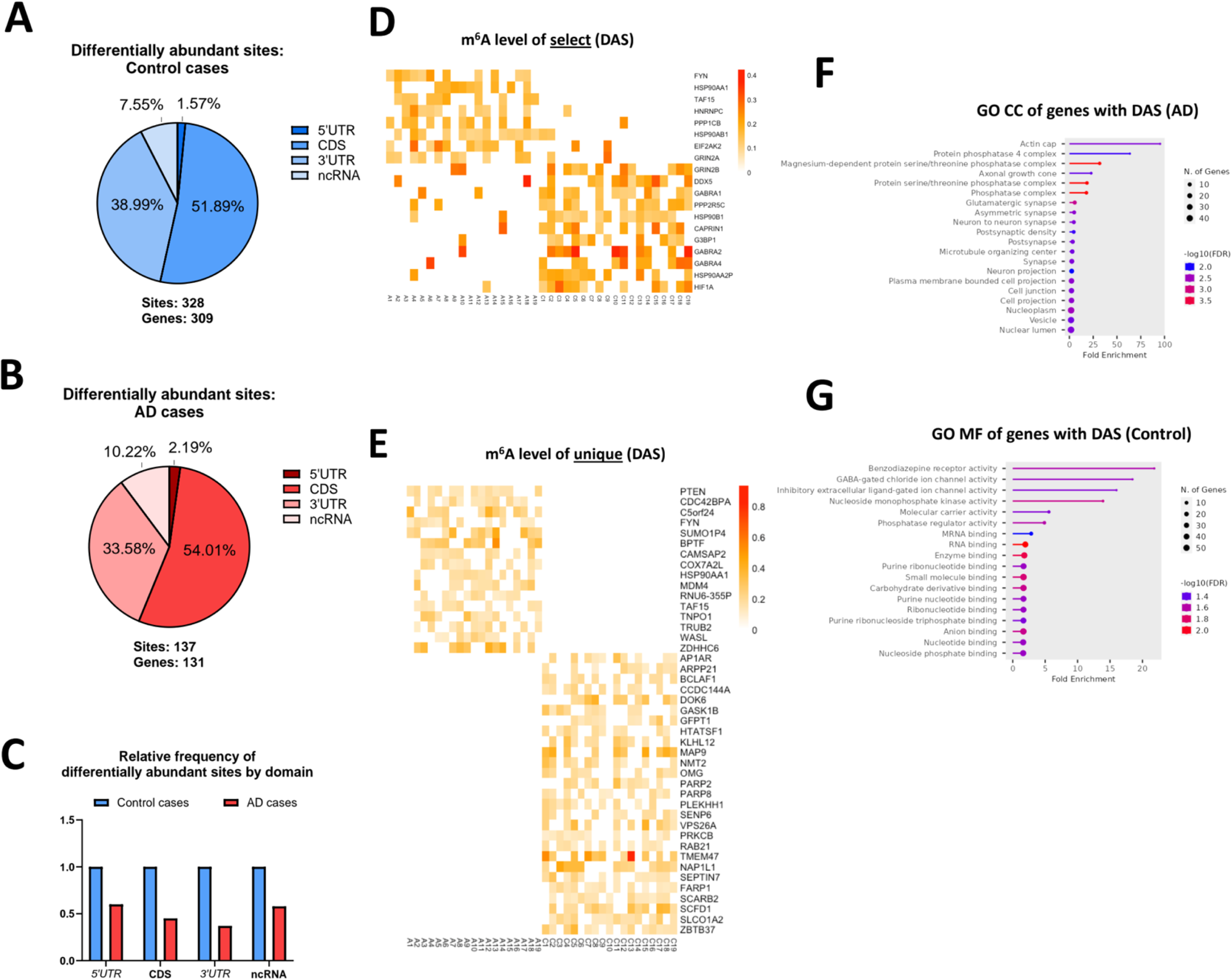
Differentially methylated site (DMS) characterization by domain distribution and gene ontology enrichment. (A) Pie chart showing percentage of Control DAS sites along different mRNA domains and on ncRNA. (B) Pie chart showing percentage of AD DAS sites along different mRNA domains and on ncRNA. (C) Bar plot showing distribution of DAS sites by mRNA domains and on ncRNA. (D) Heat map depicting m6A level (editing ratio) of select DAS among individual AD cases (A1-A19, x- axis label) and Control cases (C1-C19, x-axis label). The y-axis label is the gene that the DAS is present on. m^6^A levels (editing ratios) less than 0.1 are automatically set to 0.0 as per Bullseye parameters. (E) Heat map depicting m^6^A level (editing ratio) of unique DAS among individual AD cases (A1-A19, x-axis label) and Control cases (C1-C19, x-axis label). The y-axis label is the gene that the DAS is present on. m6A levels (editing ratios) less than 0.1 are automatically set to 0.0 as per Bullseye parameters (PMID). (F) GO CC analysis of genes with DAS on AD cases. (G) GO Molecular Function (MF) analysis of genes with DAS on Control cases.

**Supplemental Figure 9:**
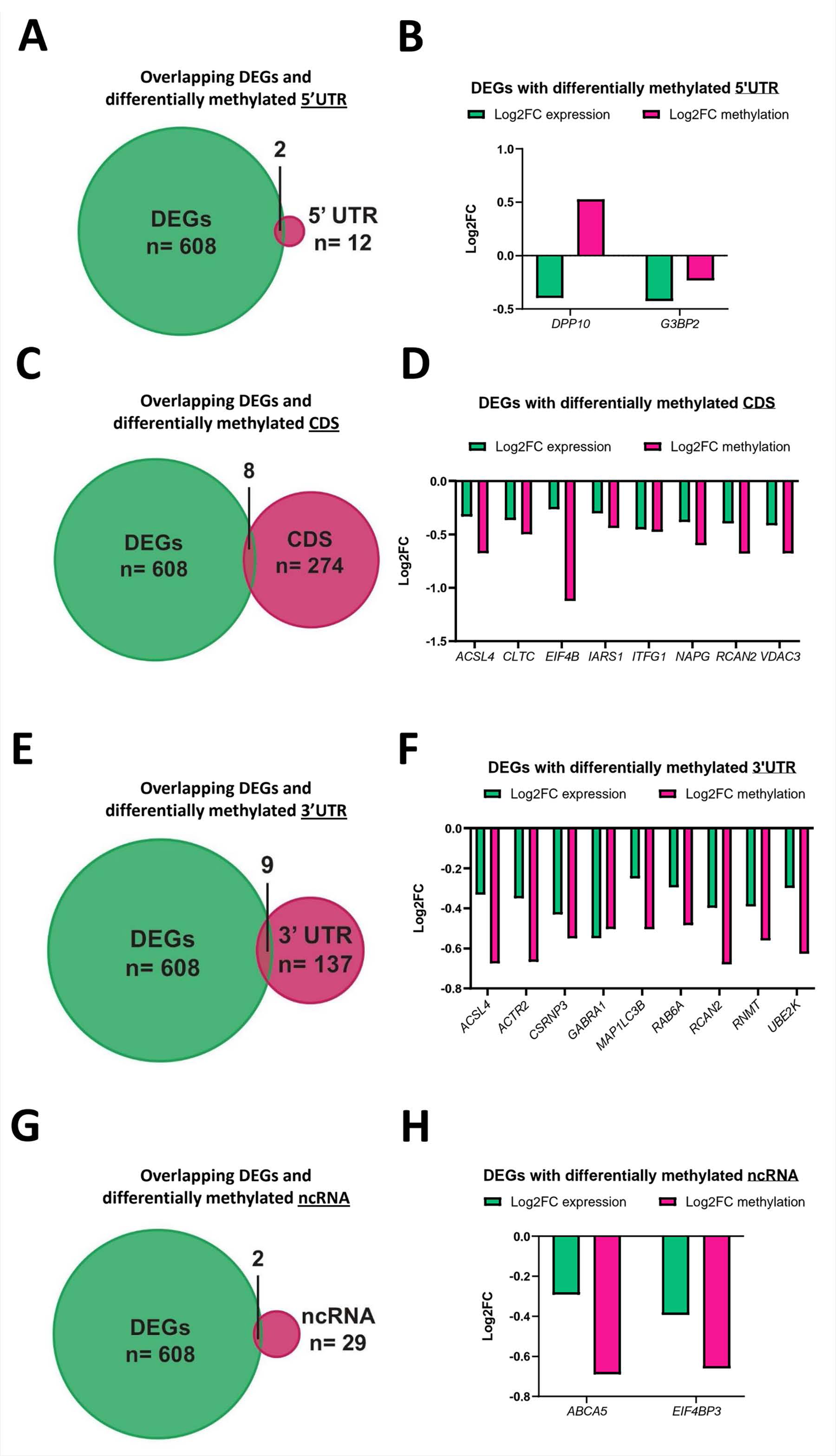
Overlapping RNA-seq and DART-seq datasets reveals shared differentially expressed genes (DEGs) with differentially methylated domains. (A) Euler plot showing the number of DEGs (n=608) and differentially methylated 5‘UTR (n=12) and genes that are both differentially expressed and differentially methylated in the 5’UTR (n=2)_ (B) Names (x-axis) and log2 fold changes of methylation and expression (y-axis) of overlapping DEGs and differentially methylated 5’UTR (C) Euler plot showing the number of DEGs (n=608) and differentially methylated CDS (n=274) and genes that are both differentially expressed and differentially methylated in the CDS (n=8). (D) Names (x-axis) and log2 fold changes of methylation and expression (y-axis) of overlapping DEGs and differentially methylated CDS. (E) Euler plot showing the number of DEGs (n=6O8) and differentially methylated 3’UTR (n=137) and genes that are both differentially expressed and differentially methylated in the 3’UTR (n=9). (F) Names (x-axis) and log2 fold changes of methylation and expression (y-axis) of overlapping DEGs and differentially methylated 3’UTR G, Euler plot showing the number of DEGs (n=608) and differentially methylated ncRNA (n=29) and genes that are both differentially expressed and have differentially methylated ncRNA (n=2). (H) Names (x-axis) and log2 fold changes of methylation and expression (y-axis) of overlapping DEGs and differentially methylated ncRNA.

**Supplemental Figure 10:**
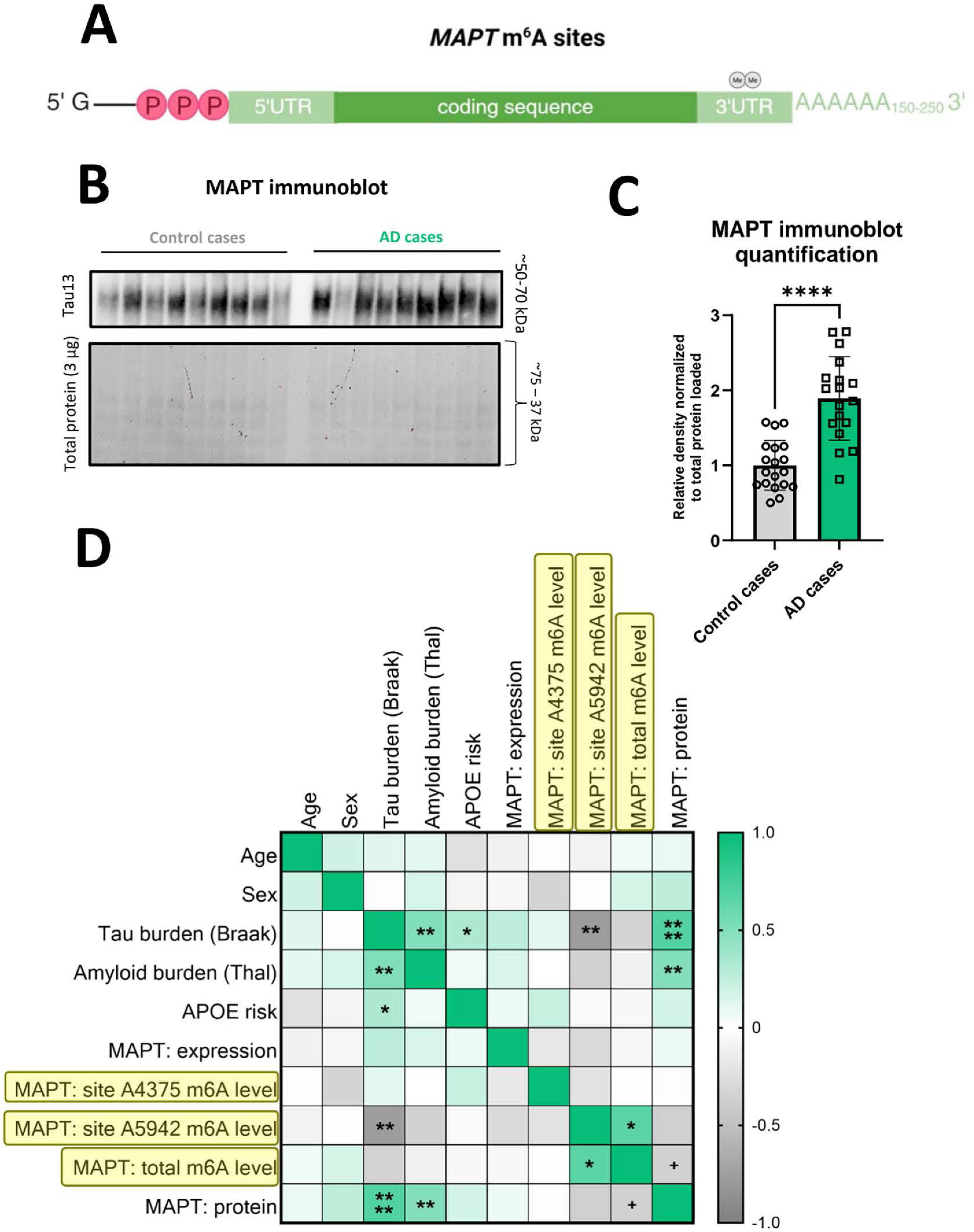
Globally hypomethylated transcripts in AD include MAPT whose m6A level associates negatively with Braak stage. **(A)** Schematics *of MAPT* transcript depicting the number and location of m6A sites identified by DART-seq. There are 2 m6A sites in the 3’UTR of MAPT, as represented by gray “Me” circles. Total sites are listed in Table S5. (B) Immunoblot comparing MAPT protein level (top) between Control and AD cases. Total protein scan (below) was used to normalize for protein input. Uncropped immunoblots can be found in Table S12. (C) Quantification of MAPT protein levels and comparison between Control and AD cases. (**** < 0.0001) P value by unpaired Student’s t test. Error bars represent mean≈ standard deviation. (D) Correlation matrix of tissue sample and case demographic information with MAPT expression, methylation, and protein levels. Pearson correlation coefficients range from -1.0 to 1.0. + P< 0.10, * P< 0.05, ** P <0.01.

**Supplemental Figure 11:**
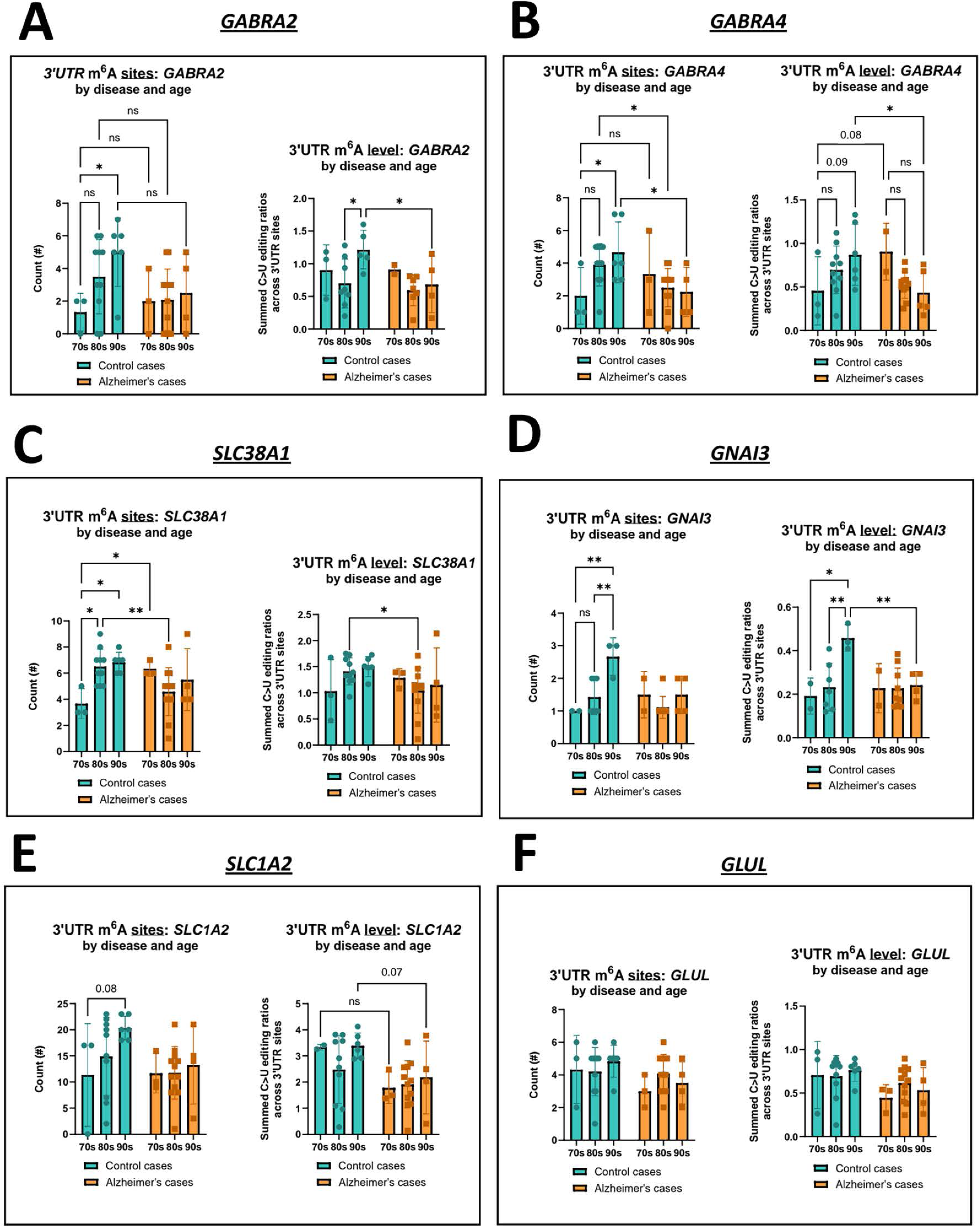
Comparison of 3’UTR-localized m*A sites and levels by disease and age of select tripartite synapse genes. (A) Comparison of 3’UTR-localized m^6^A sites (left) and levels (right) quantification of *GABRA2* between Control and AD cases grouped by age decade (70s. 80s, 90s). (* < 0.05) *P* values were generated by two-way ANOVA with Tukey’ s HSD post-hoc test. Error bars represent mean ± standard deviation (B) Comparison of 3 ‘UTR-localized m^6^A sites (left) and levels (right) quantification of *GABRA4* between Control and AD cases grouped by age. (* < 0.05) *P* values were generated by two-way ANOVA with Tukey’s HSD post-hoc test. Error bars represent mean = standard deviation (C) Comparison of 3’UTR-localized m^6^A sites (left) and levels (right) quantification of *SLC38A1* between Control and AD cases grouped by age. (* < 0.05, ** < 0.01) *P* values were generated by two-way ANOVA with Tukey’s HSD post-hoc test. Error bars represent mean± standard deviation. (D) Comparison of 3’UTR-localized m^6^A sites (left) and lewis (right) quantification of *GNAI3* between Control and AD cases grouped by age. (* < 0.05, ** < 0.01) *P* values were generated by two-way ANOVA with Tukey’s HSD post-hoc test. Error bars represent mean± standard deviation. (E) Comparison of 3’UTR-localized m^6^A sites (left) and levels (right) quantification of *SLC1A2* between Control and AD cases grouped by age. *P* values were generated by two-way ANOVA with Tukey’s HSD post-hoc test. Error bars represent mean ± standard deviation (F) Comparison of 3’UTR-localizedm^6^A sites (left) and levels (right) quantification of *GLUL* between Control and AD cases grouped by age. *P* tallies were generated by two-way ANOVA with Tukey’s HSD post-hoc test. Error bars represent mean = standard deviation

**Supplemental Figure 12:**
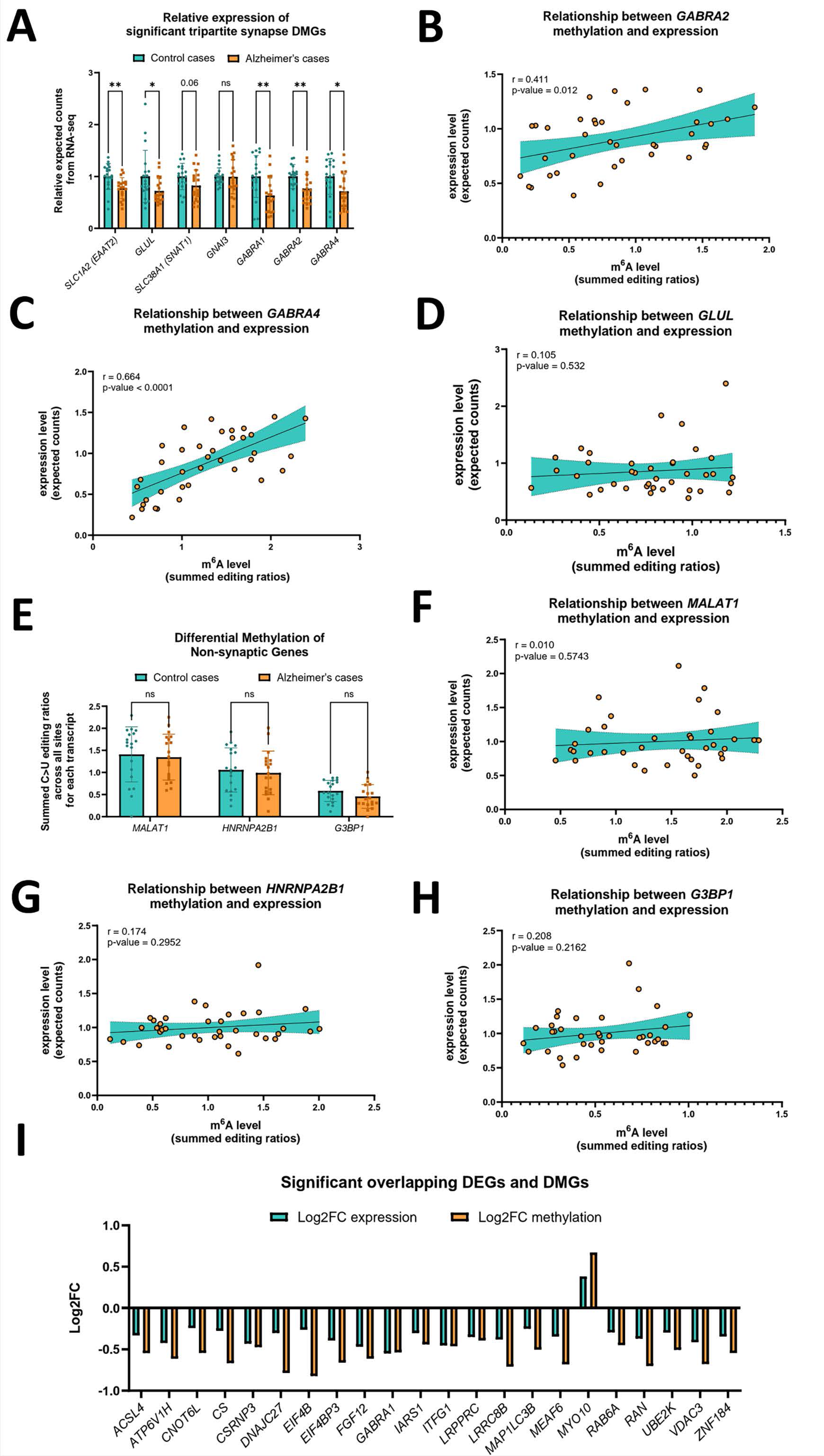
Identifying associations between transcript methylation and expression levels. (A) Comparison of relative expected counts from RNA-seq of significant differentially methylated genes (DMGs) part of tripartite synapses. *P =* 0.0850 (* < 0.05, ** < 0.01) by Mann-Whitney U test, ns = not significant. Error bars represent mean ± standard deviation. (B) Correlation of *GABRA2* m^6^A level (summed editing ratios from DART-seq) and expression level (expected counts from RNA-seq) using all brain samples. Regression is fitted to the equation: Y = 0.2263*X + 0.7029, has a positive Pearson correlation coefficient of 0.411, and is statistically significant *(P* = 0.0115). (C) Correlation of *GABRA4* m‘A level (summed editing ratios from DART-seq) and expression level (expected counts from RNA-seq) using all brain samples Regression is fitted to the equation: Y = 0.4388*X * 0.3215, has a positive Pearson correlation coefficient of 0.664, and is statistically significant (P c 0.0001). (D) Correlation of *GLUL* level (summed editing ratios from DART-seq) and expression level (expected counts from RNA-seq) using all brain samples. Regression is fitted to the equation: Y = 0.1497*X * 0.7442, has a positive Pearson correlation coefficient of 0.105, and is not statistically significant *(P* = 0.5323). (E) Comparison of m^6^A levels of non-synaptic genes between Control and AD cases. (* < 0.05) *P* values by unpaired Student’s *t* test, ns = not significant. Error bars represent mean ± standard deviation. (F) Correlation *of MALAT1* m^6^A level (summed editing ratios from DART-seq) and expression level (expected counts from RNA-seq) using all brain samples. Regression is fitted to the equation: Y = 0.06362*X + 0.9331, has a positive Pearson correlation coefficient of 0.010, and is not statistically significant *(P* = 0.5743). (G) Correlation of *HNRNPA2B1* m‘A level (summed editing ratios from DART-seq) and expression level (expected counts from RNA-seq) using all brain samples. Regression is fitted to the equation: Y = 0.08242*X + 0.9157, has a positive Pearson correlation coefficient of 0.174, and is not statistically significant (P = 0.2952). (H) Correlation of *G3BP1* m^6^A level (summed editing ratios from DART-seq) and expression level (expected counts from RNA-seq) using all brain samples Regression is fitted to the equation: Y = 0.2397*X + 0.8754, has a positive Pearson correlation coefficient of 0.208, and is not statistically significant (P = 0.2162). (I) Log2FC values of the 22 genes that were both significantly differentially expressed (p-adjusted) and differentially methylated (p-unadjusted) from the RNA-seq and DART-seq analyses, respectively. Gene names are on the x-axis. DEGs = differentially expressed genes. DMGs = differentially methylated genes.

